# Massively parallel functional testing of *MSH2* missense variants conferring Lynch Syndrome risk

**DOI:** 10.1101/2020.06.03.133017

**Authors:** Xiaoyan Jia, Bala Bharathi Burugula, Victor Chen, Rosemary M. Lemons, Sajini Jayakody, Mariam Maksutova, Jacob O. Kitzman

## Abstract

The lack of functional evidence for the majority of missense variants limits their clinical interpretability, and poses a key barrier to the broad utility of carrier screening. In Lynch Syndrome (LS), one of the most highly prevalent cancer syndromes, nearly 90% of clinically observed missense variants are deemed “variants of uncertain significance” (VUS). To systematically resolve their functional status, we performed a massively parallel screen in human cells to identify loss-of-function missense variants in the key DNA mismatch repair factor *MSH2*. The resulting functional effect map is substantially complete, covering 94% of the 17,746 possible variants, and is highly concordant (96%) with existing functional data and expert clinicians’ interpretations. The large majority (89%) of missense variants were functionally neutral, perhaps unexpectedly in light of its evolutionary conservation. These data provide ready-to-use functional evidence to resolve the ∼1,300 extant missense VUSs in *MSH2*, and may facilitate the prospective classification of newly discovered variants in the clinic.

## Introduction

Lynch Syndrome (OMIM:120435), the first discovered familial cancer syndrome^1^, confers predisposition to colorectal and endometrial cancers due to loss of DNA mismatch repair (MMR) and the resulting mutagenic burden^2–5^. Most affected individuals inherit a single loss-of-function variant in one of four MMR factors (*MSH2, MLH1, MSH6, PMS2*), followed by a somatic ‘second hit’ inactivating the remaining functional copy. Due to high prevalence in the general population (≥1:300)^6,7^, clear genetic etiology, and potential for prevention through intensified surveillance, pathogenic MMR gene variants are considered clinically actionable^8^. Though screening for MMR gene variants is increasingly routine, accurately interpreting them imposes a substantial clinical burden.

Missense changes together comprise 20-30% of Lynch Syndrome variants^9^, and are particularly challenging to interpret: their functional impacts may range from minimal to profound, and as most are individually rare, prior evidence to guide their classification is frequently unavailable. Compounding this, variants which inactivate MMR exhibit incomplete penetrance, conferring a lifetime cancer risk between 15% and 60% depending upon which gene is inactivated^10,11^. Consequently, MMR gene missense variants are not readily ruled in or out as casual, even when observed in an individual with a relevant cancer diagnosis. Indeed, the overwhelming majority of Lynch Syndrome gene missense variants listed in ClinVar (4,762/5,473, 87.0%) are deemed “variants of uncertain significance”, or VUS (Figure S1), and cannot be used to guide diagnosis and management.

Functional studies can provide a key source of evidence to support variant classification^12,13^. Traditionally, these studies are conducted *post hoc* and cannot be completed within the timeframe needed to support real-time interpretation. The heterogeneity of the functional data accumulated in the literature for MMR gene variants poses a further barrier, as the model systems and assays in use vary widely. Some assays interrogate specific mechanistic details (e.g., ATPase activity), while others report the end result upon *de novo* mutation rate. Human MMR variants have been extensively modeled in budding yeast^14,15^, but not every human MMR factor complements its yeast ortholog^16^, and many human variants occur outside of well-conserved domains and so are not readily modeled on diverged orthologs. Reconstituting MMR *in vitro* enables mechanistic studies, but may not reflect functional state under physiological conditions. Thus, while individual functional reports on MMR gene variants abound, it remains a challenge to bring this disparate body of evidence to bear on the task of classifying variant pathogenicity.

To address these challenges, we applied deep mutational scanning^17^ to prospectively and systematically measure the functional impact of missense variants in the major Lynch Syndrome gene *MSH2*. We developed human cell line models in which *MSH2* deletion was complemented by libraries of variants comprising nearly every possible *MSH2* missense allele. We leveraged an established chemical selection for mismatch repair dysfunction^18–21^, and used deep sequencing to identify the surviving *MSH2* variants. The resulting missense loss-of-function (LOF) scores are highly concordant with existing functional data and substantially outperform bioinformatic variant effect predictions. We observe that adult populations are significantly depleted for deleterious missense *MSH2* mutations, akin to what is observed for whole-gene deletions and truncating point variants in MMR genes^22,23^. These data provide a uniformly generated set of functional evidence, calibrated against known standards of pathogenicity, and will assist the interpretation of variants which would otherwise remain a source of uncertainty for clinicians, patients, and their relatives.

## Material and Methods

### *MSH2* knockout cell lines

Clonal knockout cell lines (Figure S2) were derived by transient transfection with plasmids expressing SpCas9 and guide RNAs targeting *MSH2*. For HAP1, a single guide sequence (primers JBW0001/2) targeting *MSH2* exon 6 was cloned into pSpCas9(BB)-2A-GFP (PX458, Addgene #48138) as described^24^. For HEK293, four guides (primers VC0019/20, VC0021/22, VC0023/24, VC0027/28), two flanking each side of the *MSH2* locus, were each likewise cloned into pSpCas9(BB)-2A-PuroV2 (PX459, Addgene #62988)^24^ and used in combination. Wild-type HAP1 or HEK293 cells were transfected with the respective plasmid(s) using Lipofectamine 3000 (Thermo). Clonal HAP1 *MSH2* knockout cells were isolated by sorting single GFP positive cells using a FACS Aria II instrument (BD), and colonies were screened for *MSH2* exon 6 frameshift mutations. Clonal HEK293 *MSH2* knockout cells were isolated by dilution, expanded, and screened by PCR for the presence of a full-gene knockout deletion junction(s), and the absence of an internal target in exon 12.

### *MSH2* cloning

Wild-type *MSH2* cDNA was PCR amplified from a lentiORF plasmid (Dharmacon OHS5897), and fused in-frame by HiFi Assembly (NEB) to a blasticidin resistance marker amplified from pLentiCas9Blast (AddGene #52962), separated by a P2A self-cleaving peptide linker (Figure S3). The p.Ala636Pro mutation was introduced by site-directed mutagenesis. Each resulting construct was subcloned a doxycycline-inducible lentiviral expression plasmid (pCW57.1, Addgene #41393) by digestion with SalI and NheI and ligation with T4 DNA ligase (NEB).

### Western blotting

Cells were seeded on 6-well plates, grown to confluence, and lysed by the addition of RIPA buffer and protease inhibitor cocktail (Sigma-Aldrich). Protein extracts were run for 30 min at 200 V on denaturing Bolt 4-12% Bis-Tris Plus Gels in 1× Bolt MOPS SDS buffer (Invitrogen), transferred to a nitrocellulose membrane for 1.5 h at 10 V, and blocked at 4°C overnight in 1X TBST (20 mM Tris base, 0.15 M NaCl, 0.1% Tween-20, 2% milk). Blots were probed with a 1:2000 dilution of anti-MSH2 (FE11, Thermo) and as loading control, 1:5000 dilution of anti-beta actin (BA3R, Invitrogen), followed by a 1:1000 dilution of HRP-conjugated goat anti-mouse IgG (H+L) secondary antibody (#62-6520, Invitrogen). Detection was performed with SuperSignal West Pico Chemiluminescent substrate (Thermo) and ChemiDoc XRS system (BioRad).

### Cell culture

Human HAP1 cells (Horizon Discovery) were cultured in IMDM and HEK293 and 293T/17 cells (ATCC) were cultured in DMEM, each supplemented with 10% fetal bovine serum, 100 U/ml penicillin and 100 g/ml streptomycin. During routine passaging, cells were washed with PBS pH 7.4 and dissociated with TrypLE Express. Cell lines were tested monthly for mycoplasma; all were negative.

### Library mutagenesis

*MSH2* cDNA was divided into 21 tiles (Table S1), and each was subjected to programmed allelic series mutagenesis^25^ modified as follows. Mutagenic PCR reactions used PrimeStar GXL polymerase (Takara), and 45mer mutagenesis primers (Table S2) were individually synthesized. Each codon was randomized in an individual PCR (program MUT_1, Table S3), with a common forward primer (JOK_0184) upstream of *MSH2*, and a codon-specific degenerate (NNN) reverse primer. Resulting partial-length mutant cDNA was subjected to 10 cycles of strand extension (program MUT_2) with limiting template (250 pg) of wild-type cDNA plasmid. Finally, each full-length mutant cDNA was PCR amplified with flanking outer primers (program MUT_3, primers JOK_0184 and JOK_0185). Per-codon products were pooled within each tile and purified by SPRI bead cleanup^26^. PCR products were digested with DpnI, to deplete starting plasmid, and NheI-HF and SalI-HF, for cloning into the lentiviral vector as described above. Ligation products were transformed into Endura electrocompetent *E. coli* (Lucigen). A small aliquot of the transformant culture was plated to monitor library complexity, while the remainder was expanded for 12-16 hours at 32°C in Luria Broth with 100 ug/ml ampicillin (LB+amp). We required total colony forming units ≥ 4×10^4^, for mean ∼10-fold coverage per variant. Lentiviral tile plasmid libraries were isolated using ZymoPURE II Plasmid kit, and converted to shotgun libraries for deep sequencing by Tn5 tagmentation^27^, to assess mutational specificity and coverage (Figure S4). Absent or under-represented codons were individually retrieved from the earlier mutagenesis steps, pooled, cloned as above, and spiked into each tile library as needed to equalize coverage. Finally, these mutant tile libraries, as well as control clones (WT and A636P), were modified to add trackable barcodes. Each clone or library was linearized with SalI-HF (NEB) downstream of the *MSH2*-2A-bl^R^ ORF, and a degenerate 20-mer (primer jklab0172) was inserted by HiFi Assembly (NEB). The resulting barcoded clone pools were transformed, expanded, and isolated as described above.

### Lentiviral preparation and transduction

Barcoded *MSH2-*2A-bl^R^ cDNA libraries were packaged into lentivirus by co-transfecting HEK293T/17 cells (ATCC) with the transfer plasmid pool plus envelope and packaging vectors (pMD2.G, Addgene #12259 and psPAX2, #12260). For each pool, 4.4×10^6^ cells were plated in a 100 mm dish, then transfected with 17 ug total plasmid (2.0:1.0:1.3 transfer:envelope:packaging ratio), using Lipofectamine 3000 (Thermo). Media was replaced at 6 h post transfection, and viral supernatants were collected at 24 h, passed through an 0.45 micron filter (Millipore), and used immediately or stored at −80°C until use. Viral titer was estimated by transduction with a dilution series followed by puromycin selection and CellTiterGlo (Promega) cell proliferation assay^28^, and verified by counting unique barcodes in the transduced population (barc-seq, described below). For each tile, *MSH2* knockout HAP1 or HEK293 cells were transduced with mutant library at low multiplicity of infection (< 0.1), by applying 1.5 ml viral supernatant (with 8 µg mL^-1^ polybrene) to each of four 100mm dishes (Corning) containing ∼7.5×10^6^ cells. After 48 h, transduced cells were selected by addition of 1 µg mL^-1^ puromycin, which was included in all subsequent steps.

### Mismatch repair functional selection

Cells expressing loss-of-function *MSH2* variants were enriched by adding 6-thioguanine (6-TG) to the culture media. To find an optimal concentration, parental and *MSH2* KO cell lines were plated in quadruplicate wells of a 96-well plate and treated with a range of 6-TG concentrations. Cell growth after 4 days was measured using the CellTiterGlo 2 kit (Promega) and a GloMax luminometer (Promega), and survival was normalized to untreated wells (Figure S2).

For library selections, each mutant tile cell population was spiked with barcoded knock-in control cells (*MSH2* WT: to 10% of the population, A636P: to 0.5%), then split and expanded in triplicate 10 cm dishes (Figure S5-S6) in media supplemented with 1 µg mL^-1^ doxycycline to induce *MSH2* expression. From each baseline passage (“P0”) culture, 10^7^ cells were collected for gDNA extraction, and cells were split into 6-TG treated and mock-treated cultures, with media containing 1 µg mL^-1^ puromycin, 1 µg mL^-1^ doxycycline, 5 µg mL^-1^ blasticidin (to enforce *MSH2* expression), with and without 1 uM 6-TG, respectively. Cells were grown to confluence for two rounds, at each round harvesting ≥10^7^ cells for gDNA extraction and seeding with 3×10^6^ cells to avoid population bottlenecks.

### Amplification and sequencing of integrated *MSH2*

After selections, genomic DNA was isolated from 1-3×10^7^ cells using the Quick-DNA Midiprep Plus kit, and sequenced in tiles. To provide sufficiently redundant sampling of integrated *MSH2* libraries, for each sample, 16 replicate PCR reactions were assembled, each with 0.5 µg of gDNA as template (template copy number of 2.4×10^6^ genome equivalents per tile). Integrated inserts (∼3.7 kbp), including *MSH2* cDNA and associated barcode, were PCR amplified with PrimeSTAR GXL (program MSH2gDNA). Each set of 16 replicate amplicons was pooled and purified with SPRI beads. Next, tile-seq libraries were prepared from each pool of amplified integrated *MSH2* constructs. Tile-specific primers (Table S2) were used to further amplify the tile mutagenized in each sample in a PCR reaction (program TILESEQ) with 10 ng full-length amplicon (≥ 2.6×10^9^ copies) as template. Dual-indexed illumina adapters (sequences available upon request) were then added by a subsequent round of PCR. The resulting short amplicon libraries thus provide overlapping read coverage for most of each tile. Tile-seq libraries were pooled for paired end 150bp Illumina sequencing (≥ 1×10^6^ paired reads per sample, n=189 libraries). Pool complexity (the number of distinct transduced clones) was assessed by tile-seq (Figure S7) and barc-seq (Figure S8). Barcodes were amplified from the full-length *MSH2* amplicon and converted to illumina amplicon libraries, as for tile-seq, but using primers jklab0077 and jklab0078 (program BCSEQ). Each tile’s starting population (“P0”) had ≥20,000 barcodes, indicating at least that number of unique initial transduction events. Catalogs of barcodes associated with each of the two *MSH2* control clones (WT, A636P), and for each tile library, were established by barc-seq (as described above) from their respective gDNAs at P0 (after transduction). Finally, tile-seq libraries were prepared from a fully wild-type *MSH2* clone plasmid DNA, to measure and correct position-specific errors, primarily single base substitutions which varied considerably by position and type (Figure S9)

### Sequence data processing

Shotgun sequencing reads were aligned to a set of *MSH2* cDNA sequences with each codon sequentially replaced with “NNN” using bbmap (https://jgi.doe.gov/data-and-tools/bbtools/) in semi-perfect mode, to identify reads containing at most one mutated codon with between one and three edits. Custom python scripts converted the resulting alignments to a per-sample table of counts for every possible codon mutation. For tile-seq libraries, paired reads were overlapped and error-corrected with PEAR v0.9.6^29^; only perfectly matching pairs were used. For each tile, all possible single-codon mutation sequences were enumerated, and the count of reads exactly matching each mutant sequence was tabulated. Reads with additional mutations or sequencing errors were discarded. Barc-seq reads were clustered with starcode v1.3^30^ to define and count unique barcodes. To determine the number of distinct barcodes present, robust to sequencing or amplification artifacts, we ranked clustered barcode groups by read count and determined the minimum number of barcodes for which all equally or more abundant barcodes cumulatively accounted for ≥90% of the total read counts.

### Variant enrichment scoring

The raw frequency of each per codon mutation *i* in each sample *s* was calculated, with a pseudocount of 0.9: 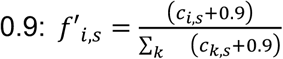. Position specific error rates from wild-type *MSH2* plasmid tile-seq were subtracted and normalized to yield corrected frequencies: 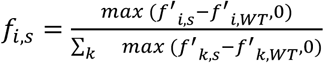. Variants with frequency ≤1/200,000 in the initial transduced cell population (P0) were removed from further analysis. Next, variants substantially enriched or depleted after mock treatment (absolute log2 ratio > 2, after two passages blasticidin vs P0) were likewise removed; these variants may be *in cis* with a frameshift or premature truncating mutation beyond the sequenced tile. A loss-of-function (LOF) score was taken as the log2ratio of the frequency after 6-TG treatment over that after mock treatment. For amino acid substitutions represented by more than one equivalent codon substitution, the median was taken across those codons’ LOF scores to yield an amino acid-level LOF score. Within each replicate, variants were removed when represented by exactly two LOF scores with opposite signs. Finally, the median score was taken for each amino acid LOF score across the three replicates; when only two replicate scores remained and had discordant signs, they were removed. False discovery rate was estimated by random sampling of synonymous variants’ read counts, with replacement and keeping triplicates together. To mimic the number of codons corresponding to a given mutant amino acid, synonymous variants were sampled in groups of size one to six. Counts were processed to LOF scores and filtered as described above, combining first within each simulated missense amino acid and then across replicates; final LOF scores greater than zero were taken as false discoveries. Overall FDR was estimated as the mean FDR for amino acids with each number of different variant codons (1, 2, 3, 4, or 6 codons), weighted by the fraction of variant amino acids with that number of codons.

### Clinical and population variants

*MSH2* missense variants and classifications were obtained from ClinVar (http://www.ncbi.nlm.nih.gov/clinvar/)” www.ncbi.nlm.nih.gov/clinvar/) on 2020 April 14. Redundant variants were removed, and likely splice-disruptive variants with a SpliceAI^31^ score of ≥0.2 were filtered out, along with those where the ClinVar record indicated a known splice-disruptive mechanism (Table S4). Only variants with expert panel reviews were included in comparisons to LOF scores. *MSH2* missense variants and classifications were accessed from InSiGHT (http://www.insight-group.org/variants/databases/)” www.insight-group.org/variants/databases/). *MSH2* missense population variation was obtained from the gnomAD database^23^, combining whole-genome and exome calls from versions 2.1.1 and 3.0. The expected number of LOF missense mutations was modeled with a binomial distribution parameterized by *n*=744 (the number of scored missense variants in gnomAD) and *p* taken as the sum of LOF missense point variants’ relative likelihoods of occurring *de novo*^*32*^, equivalent to drawing sets of size 744 from the universe of possible missense SNVs with probabilities scaled by their individual relative likelihoods.

### Bioinformatic classification

All *MSH2* missense variants were scored by the following bioinformatic tools:CADD^33^, PolyPhen-2^34^, REVEL^35^, PON-MMR^36^, MAPP-MMR^37^, and FoldX (scores obtained from ^38^). For CADD and REVEL, scores were available only for missense variants reachable by single-base variants (SNVs); for amino-acid substitutions that could arise from more than one SNV, the mean of those SNVs’ scores was taken.

### MSH2 structure and conservation

The MSH2 crystal structure^39^ was obtained from the Protein Databank (accession 2O8E) and rendered with PyMOL. Amino acid secondary structure assignment was extracted from pre-calculated DSSP files^40^. Surface accessibility was calculated using ASA (http://cib.cf.ocha.ac.jp/bitool/ASA/). MSH2 ortholog protein sequences were downloaded from Ensembl (https://useast.ensembl.org/) and aligned with MUSCLE^41^.

## Results

### A Human Cell Platform to Test *MSH2* Variant Function

We established a human cell system to model *MSH2* variant function using the mismatch repair proficient^42^ cell line HAP1 (Figure 1A, C). First, to disrupt MMR, we derived clonal *MSH2* knockout cells bearing a 19-bp frameshift deletion in *MSH2* exon 6, and verified that it lacked detectable MSH2 protein expression (Figure S2). To restore MMR, we stably reintroduced into cells either wild-type *MSH2* cDNA (KO+WT) or a pathogenic founder allele (KO+A636P) on inducible lentiviral constructs. After inducing expression with doxycycline, wild-type MSH2 protein reached near-endogenous levels, while A636P was only barely detectable, consistent with the known destabilizing effect of this variant^43^ (Figure S3).

**Figure 1.**
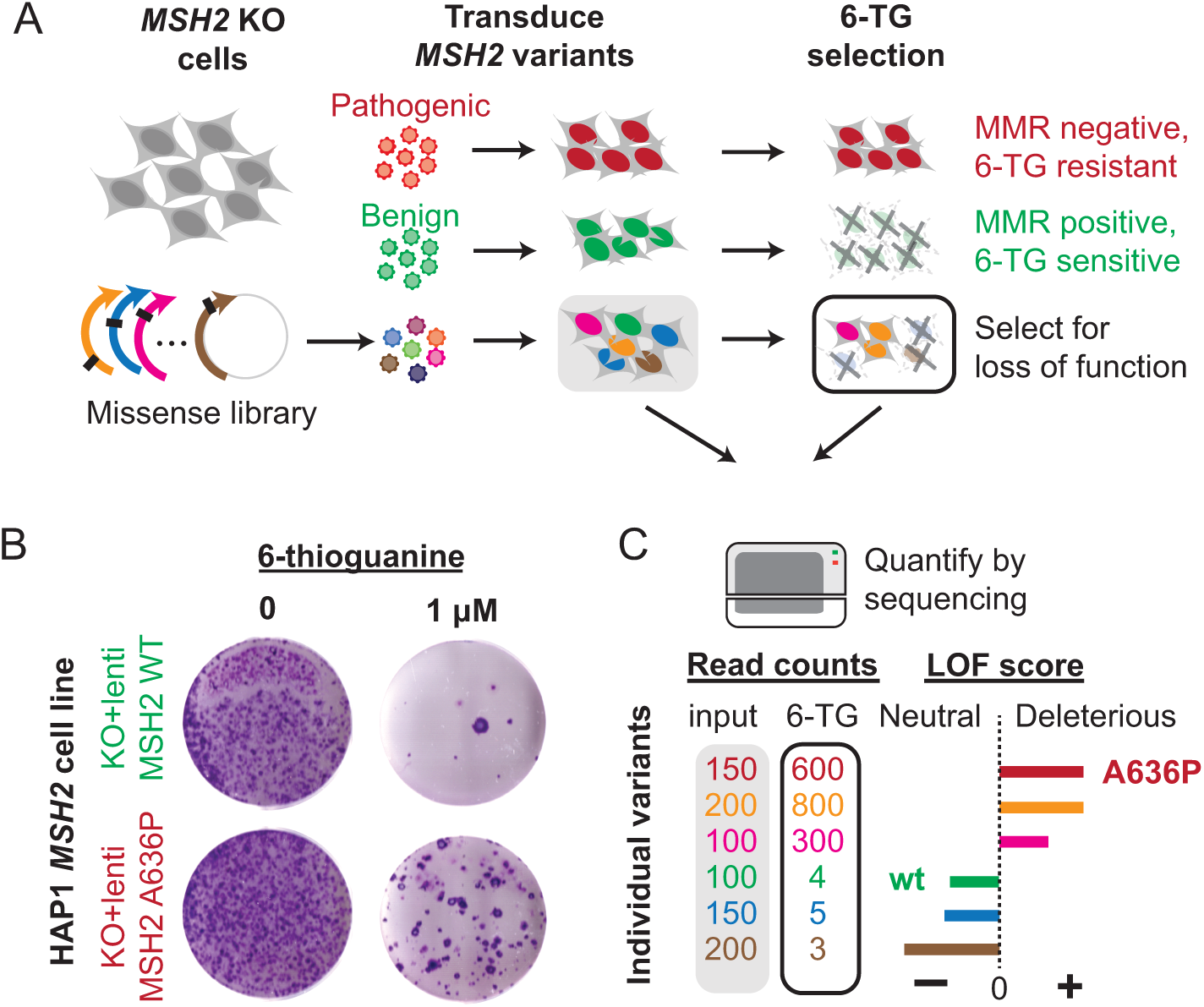
Overview of *MSH2* functional screen. (A) Generating and testing *MSH2* variant function in isogenic human cells. Cells with pathogenic loss-of-function variants (top) are resistant to 6-thioguanine (6-TG), while complementation by a benign functional variant (middle) restores 6-TG sensitivity. In a pooled screen (bottom), a mixed library of *MSH2* missense variants is transduced, and 6-TG treatment selects for loss-of-function variants. (B) 6-TG resistance reflects MMR function in human cells. HAP1 *MSH2* KO cells were transduced with either functional (wild-type) *MSH2* cDNA (upper row) or loss-of-function missense variant A636P (bottom row), and grown for 4 days without (left) or with 6-TG (1 uM, right). (C) Readout using deep sequencing to quantify abundances of each *MSH2* allele before and after 6-TG selection. The resulting loss-of-function (LOF) score is positive for deleterious variants and negative for functionally neutral ones.

As a readout for *MSH2* function, we leveraged selection with the purine analog 6-thioguanine (6-TG)^19,44^. Incorporation of 6-TG is selectively toxic to MMR-proficient cells, as it creates lesions that the MMR machinery recognizes but cannot repair, culminating in replication arrest^20,21^. As expected, KO cells showed increased 6-TG resistance compared to parental HAP1 cells, while 6-TG sensitivity was restored by reintroduction of wild-type *MSH2* but not the pathogenic A636P mutant (Figures 1B and S5). To mimic simultaneously testing thousands of mutations, we prepared a mixed culture of KO+WT cells and KO+A636P cells, with the respective *MSH2* expression constructs bearing distinct, identifying barcode libraries *in cis* (Figure S5). We quantified barcode abundances by deep sequencing (**Material and Methods**), and found barcodes linked to the pathogenic variant A636P were strongly enriched relative to WT barcodes (median fold change: 179, *P*<2.2×10^−16^, Kolmogorov-Smirnov test). This result confirms the feasibility of highly multiplexed MMR functional tests in isogenic human cell mixtures.

### A Functional Effect Map Covering >17,000 *MSH2* Missense Variants

To model known and novel *MSH2* variants, we applied single amino-acid saturation mutagenesis^25^ to generate libraries comprising every possible missense, synonymous and nonsense variant (**Material and Methods**). *MSH2* cDNA (2,802 bp) was divided into 21 tiles to enable tracking by amplicon sequencing^45^, and to mitigate the impact of recombination during lentiviral replication^46^. The mutant cDNA library for each tile was cloned into the inducible lentiviral vector mentioned above, followed by a C-terminal 2A linker and a blasticidin resistance gene, to permit selection for complete in-frame cDNA expression (Figure S3). HAP1 *MSH2* KO cells were transduced at low multiplicity (<0.1) to yield a population in which each cell expressed a single mutant *MSH2* variant. Sequencing of integrated libraries from genomic DNA confirmed that nearly all single-codon mutations were present (97.3%) with highly uniform representation (Figure S7). Sequencing errors were modeled empirically by deeply sequencing a wild-type *MSH2* plasmid, and used to correct error-prone positions (Figure S9).

Each *MSH2* mutant cell pool was then selected *en masse* for MMR deficiency. To allow comparison across pools, each was spiked with barcoded control cells (KO+WT: 10% of cells; KO+A636P: 0.5%). Pools were then grown under selective (6-TG+blasticidin) or mock (blasticidin only) conditions for two passages (Figure S6). Barc-seq of the integrated *MSH2* cDNAs confirmed that, as expected, the spiked-in *MSH2* A636P cells were consistently enriched over WT by 6-TG selection (median fold-change relative to mock treatment: 105; Figure S5).

To track the functional status of each *MSH2* variant, we performed amplicon sequencing of each tile throughout the course of selection. Loss-of-function (LOF) scores for each variant were calculated as the log2-ratio of that variant’s frequency after 6-TG treatment divided by its frequency after mock treatment, such that deleterious variants should score positively and neutral variants negatively (Figure 1C). LOF scores strongly correlated across replicate 6-TG selections (mean pairwise Pearson’s r=0.77, Figure S10). As expected, mock treatment had little effect upon most variants’ frequencies, except for truncating variants which terminate translation before the downstream blasticidin resistance marker and were therefore depleted (Figure S11).

The resulting functional effect map was substantially complete, covering 94.4% of possible *MSH2* single-codon substitutions (Figure 2). Missense variant scores were bimodal, with the large majority (89.4%) scoring negatively, suggesting a high degree of tolerance to single amino acid substitution (Figure 2G). Indeed, for over half of MSH2 residues (510/934), substitution to any other amino acid was tolerated. As expected, synonymous variants predominantly scored negatively (820/841, 97.5%), with exceptions likely caused by loss of function from an additional, non-programmed mutation outside the sequenced tile. Most of the scored missense variants were represented by multiple equivalent codons (on average, 3.0), and summarizing these internal replicates into a single, amino-acid level LOF score resulted in an overall false discovery rate of 0.95%. To verify these results were generalizable across cell types, we repeated the screen for two tiles (tile 14 and 15) in another MMR-proficient human cell line, HEK293, with largely concordant results (Figure S12).

**Figure 2.**
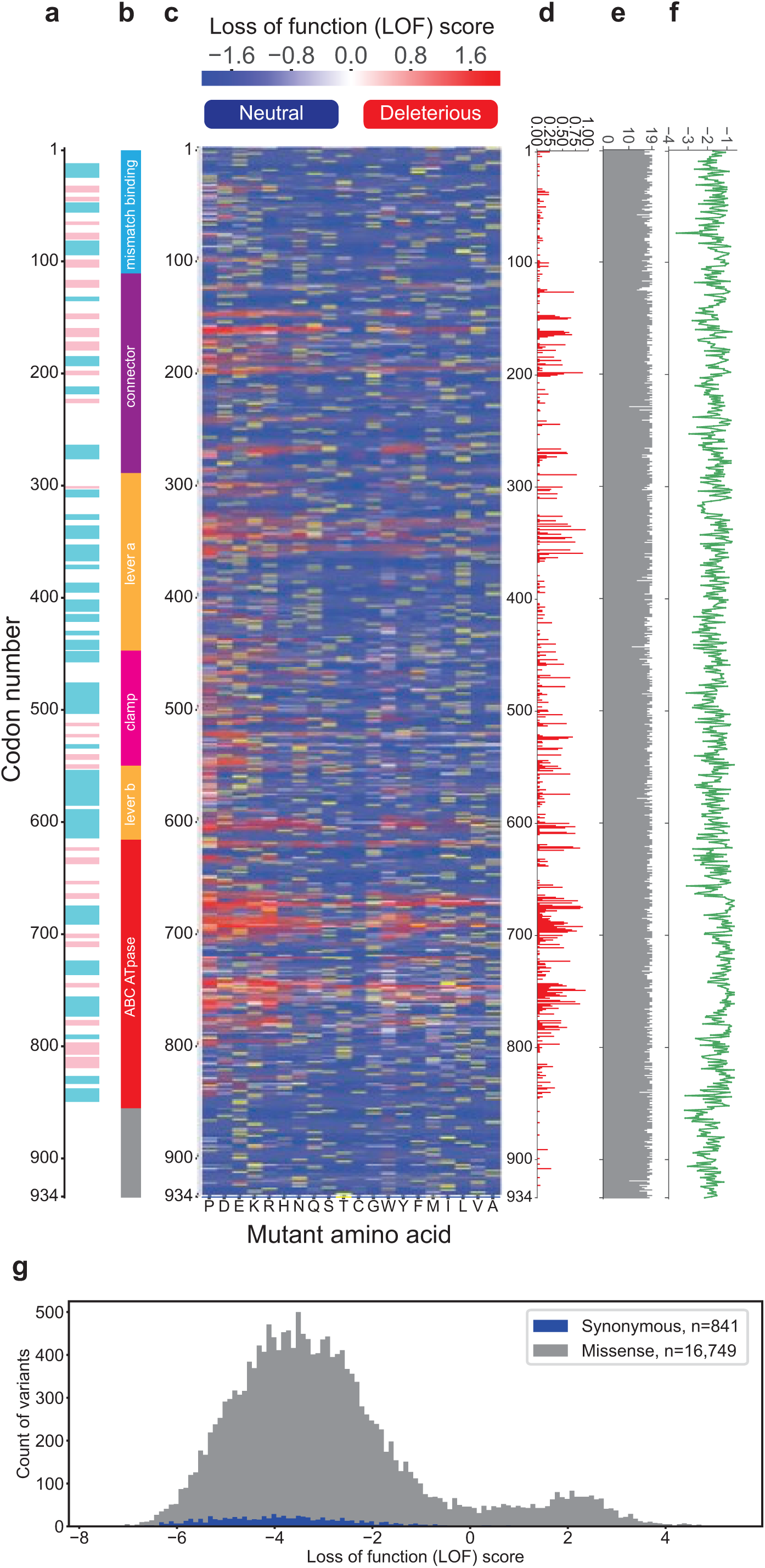
Functional effect map of *MSH2* missense variants. (A) MSH2 secondary structure (cyan: alpha helices, pink: beta sheets), and (B) protein domains. (C) Heatmap of loss-of-function (LOF) scores after mutating each of 934 residues in MSH2 (rows) to each of 19 other possible amino acids (columns). Negative scores (shaded blue) indicate functionally neutral variants, while positive scores (red) indicate deleterious ones; Yellow denotes wild-type amino acid, gray: no data. (D) Fractions of substitutions which are disruptive (LOF score>0) by position. (E) Number of scored missense variants by position (max. possible: 19). (F) Evolutionary conservation (PSIC score) by position. (G) Distributions of LOF scores, shaded by variant class.

### Pooled Measurements Recapitulate Existing Variant Interpretations

We sought to validate these pooled measurements by comparison to traditional, low-throughput functional studies. We collated a set of 184 *MSH2* missense variants previously characterized as functionally deleterious or neutral by individual cell-based^15,19,44,47–51^ and/or biochemical assays^52,53^, discarding four variants where multiple studies disagreed and eight predicted to impact splicing, an effect not measured by this approach (SpliceAI^31^ score>0.2; Tables S4-5). Our LOF scores agreed with these earlier reports for the great majority of variants covered (158/165, 95.8%; Figure 3A). We examined the variants at which our results and previous reports disagreed, and noted that for four of those seven, our LOF scores were internally consistent across multiple codons encoding the same amino acid substitution (Figure S13). These may reflect false negatives or positives on the part of the previous screens, or differences between this human cell model and previous yeast or *in vitro* assays.

**Figure 3.**
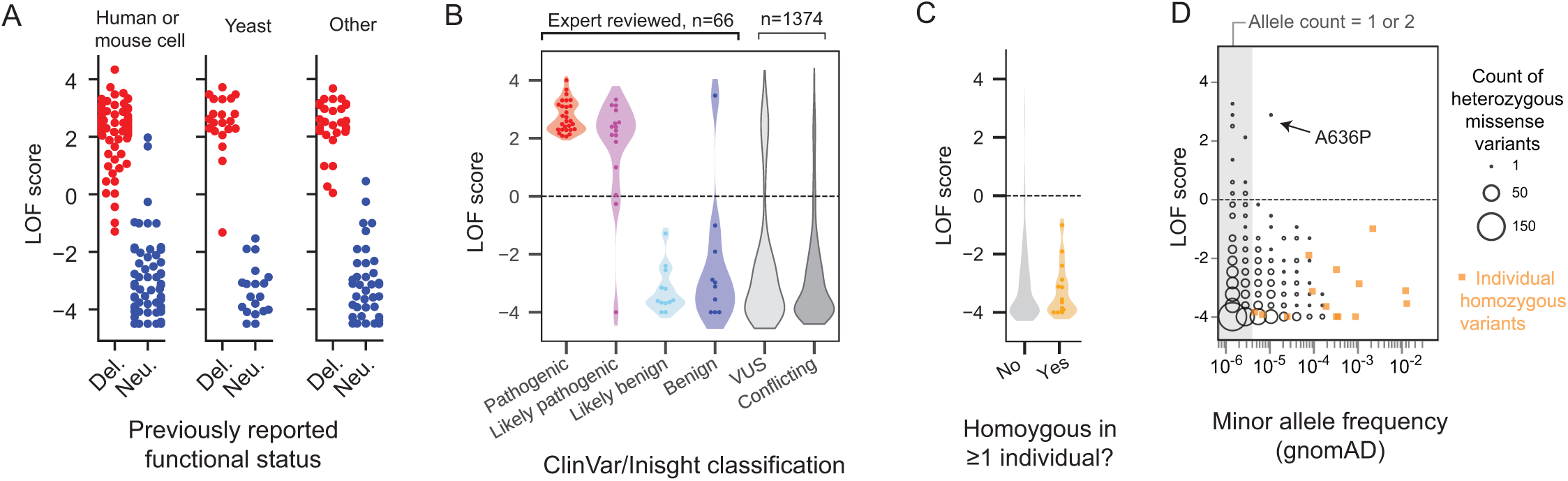
Concordance with existing functional and clinical classifications. (A) Measured LOF scores (y-axis) for published *MSH2* missense variants (n=295 reports covering 184 distinct variants), separated by previous characterization (x-axis) in individual cellular or biochemical assays. (B) LOF scores for patient missense variants from ClinVar and InSight databases, by variant classification. (C) Distributions of LOF scores of *MSH2* missense variants in the gnomAD database^23^, for variants which do not appear in the homozygous state (gray) or those which do in ≥1 individual (orange). (D) LOF score versus gnomAD allele frequency for *MSH2* missense variants; size of circle denotes number of heterozygous variants; orange squares mark missense variants observed in homozygous state in ≥1 individual. Gray region denotes singletons/doubletons (gnomAD allele count ≤2).

To assess the suitability of these scores for classifying variant pathogenicity, we intersected them with clinical variant databases. We retrieved all *MSH2* missense variants with consensus annotations from InSight^9^ or expert review in ClinVar, totaling 75 variants. After excluding nine variants predicted by SpliceAI or annotated in ClinVar as splice-disruptive, our LOF scores agreed with clinical annotations for 63 of the 66 remaining variants (95.5%, Figure 3B). These data provide evidence to facilitate interpretation of the 1,374 variants without classification (i.e., VUS) or with conflicting reports. Our scores predicted that 112 of these (8.2%) unresolved variants are functionally deleterious, similar to the overall rate of LOF among all missense variants reachable by a single nucleotide change (7.7%; *P*=0.51, two-sided binomial test). This result highlights the pervasiveness of benign missense variation in clinical databases and identifies a subset of variants likely to be causal for Lynch Syndrome.

We next examined how these function scores varied in apparently healthy populations (Figure 3C-D). The gnomAD database^23^ lists 744 of the scored *MSH2* missense variants, nearly all rare (742/744 with minor allele frequency < 0.5%). Among these, only 18 scored as deleterious in our assay (all singletons or doubletons except the founder allele A636P), a significant depletion (*P*=6.9×10^−7^, two-sided binomial test) relative to randomly drawn, size-matched sets of single-base missense variants. All 14 *MSH2* missense variants observed in the homozygous state in gnomAD scored as functionally neutral, as expected given that biallelic MMR loss causes pediatric-onset cancer syndromes. These results reflect survivorship bias which depletes adult cohorts of deleterious variants in genes underlying disorders such as Lynch Syndrome, as previously noted for *MSH2* deletions^22^.

### Loss of Function Scores Outperform Bioinformatic Predictors

Bioinformatic tools are frequently used to interpret clinical variants when other forms of evidence, such as co-segregation with disease, functional studies, or population variant allele frequency, are unavailable or uninformative^54^. We compared the classification performance of our LOF scores with that of six computational predictors, including three specific to MMR genes: MAPP-MMR^55^, FoldX^56^ thermostability predictions from Nielsen et al^38^ and PON-MMR^57^, and three general-purpose tools REVEL^35^, CADD^58^, and PolyPhen2^34^. As a truth set, we used the missense variants with previously reported functional evidence (Figure 3A). LOF scores outperformed all bioinformatic classifiers in recapitulating these functional classifications, reaching an area under the precision recall curve (auPR, Figure 4A) of 0.990, which exceeded the six algorithms’ (auPR range: 0.682-0.921). MMR-specific classifiers performed markedly better (auPR mean: 0.867) than general-purpose, pathway agnostic methods (auPR mean: 0.741), which displayed poor specificity, possibly reflecting the challenge of distinguishing functional effects between variants in highly conserved genes such as *MSH2*.

**Figure 4.**
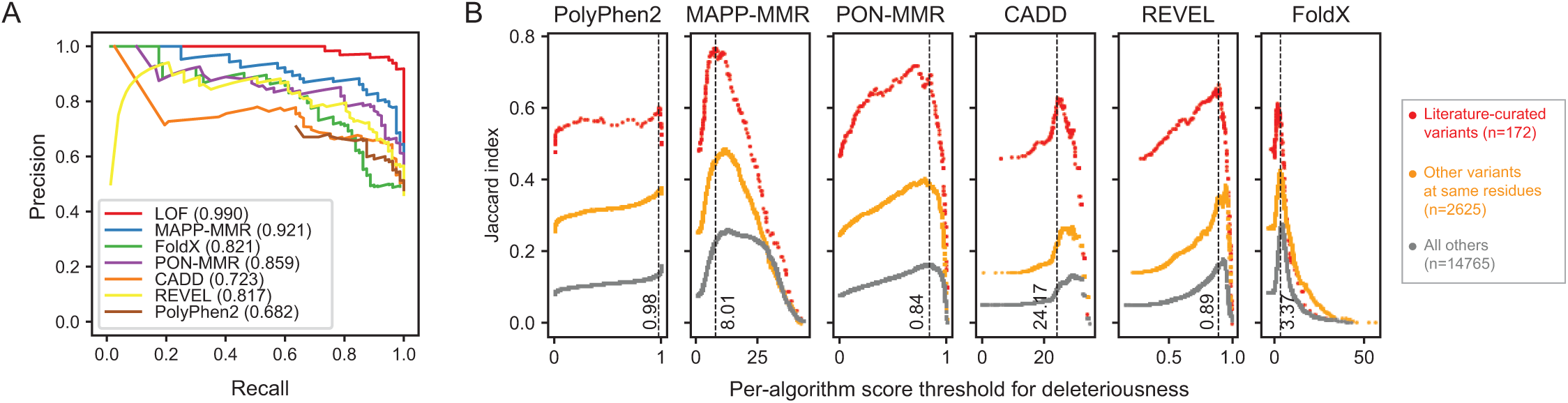
Comparison with bioinformatic predictions of pathogenicity. (A) Precision-recall curves show LOF scores outperform all tested bioinformatic callers in predicting the functional status of the validation set of variants with known effect; area under the curve indicated for each. (B) Concordance of each bioinformatic predictor (Jaccard Index) with experimental LOF scores is plotted versus algorithm-specific score threshold for deleteriousness. Algorithms were most concordant with LOF scores among mutations with published functional data (red), followed by other mutations at the same residues as those mutations (orange) and mutations all other residues (gray). Dotted line and score indicate optimal threshold for each predictor on known the validation set.

We also tested performance of the experimental and bioinformatic scores using a fixed cutoff selected for each, as would be required in practice. For LOF scores, a score cutoff of 0 was selected *a priori*, taking as deleterious any variants enriched by 6-TG. For each bioinformatic algorithm, we selected a cutoff (Figure S14A-B) which maximized the Matthews Correlation Coefficient (MCC), a balanced measure of specificity and sensitivity^59^, across the full validation set. These were broadly similar to cutoffs used previously; for instance, by an earlier estimate^60^, the CADD score cutoff of 24.17 used here corresponds to a 96.0% probability of being an Insight Class 5 (pathogenic) variant. Using these cutoffs, LOF scores reached an MCC of 0.92, higher than all six tools’ (range: 0.50-0.74; Figure S14A). Notably, because optimizing a cutoff score for each tool using the full validation set amounts to training and testing on the same data, this comparison may overestimate their performance relative to a test on newly seen alleles, whereas the LOF scores should have no such bias.

Although the bioinformatic predictions were modestly correlated with our experimental measurements for variants in the functional validation set, they were markedly less concordant for other variants at those same residues, or throughout *MSH2* at large (Figure 4B). Similarly weak overall agreement has also been observed when benchmarking bioinformatic classifiers with deep mutational scans of other genes^61–63^. As variant effect predictors are often trained on the limited number of known variants with available classifications, their divergence with our experimental measurements may reflect overfitting, further suggesting that the comparison to a small set of functionally characterized alleles overestimates bioinformatic predictors’ performance.

### Functional Constraint Highlights Structural Features

Deep mutational scans can inform protein structure-function relationships^64,65^. On aggregate, chemically conservative substitutions across MSH2 were rarely deleterious (5.63%), compared to substitutions replacing hydrophobic residues with charged ones (28.0%; Figure S15). Mapping function scores onto the crystal structure^39^ (Figure 5A), we noted several regions of mutational intolerance corresponding to structural features. The phosphate-binding Walker A motif (residues 674-676) is required for the essential ATPase activity of MSH2^66^ and was almost completely intolerant to mutation (50/55 mutations LOF, Figure 5B,C). We also noticed strong constraint at glycine 126, possibly reflecting the requirement for flexibility to accommodate conformational shifts between the mismatch binding and connector domains (Figure 5C)^39^. The connector domain’s hydrophobic core, comprising three beta sheets and inward-facing residues of the surrounding helices, was also largely functionally intolerant to substitutions. Conversely, despite its name, the N-terminal mismatch binding domain (residues 1-124) was largely devoid of missense LOF variants, consistent with its dispensability in yeast^67^, and observations that *MSH2* start codon loss is not strongly pathogenic in humans^68,69^. By contrast, prokaryotic mutS homodimerizes, with both N termini making extensive DNA contacts essential for mismatch recognition^70^. Several positively charged clamp domain residues (K512, R524, K546), which interact with the DNA backbone by hydrogen bonding^39^, were strongly intolerant to substitution, especially to negatively charged residues. Although computational predictions of protein stability^38^ showed modest overall correlation with LOF scores (Pearson’s r=0.39, Figure S14), we noted a few exceptions including the ATPase Walker motifs, which showed strong constraint in our data but were not predicted to be destabilizing (Figure S15), suggesting mechanisms other than protein instability.

**Figure 5.**
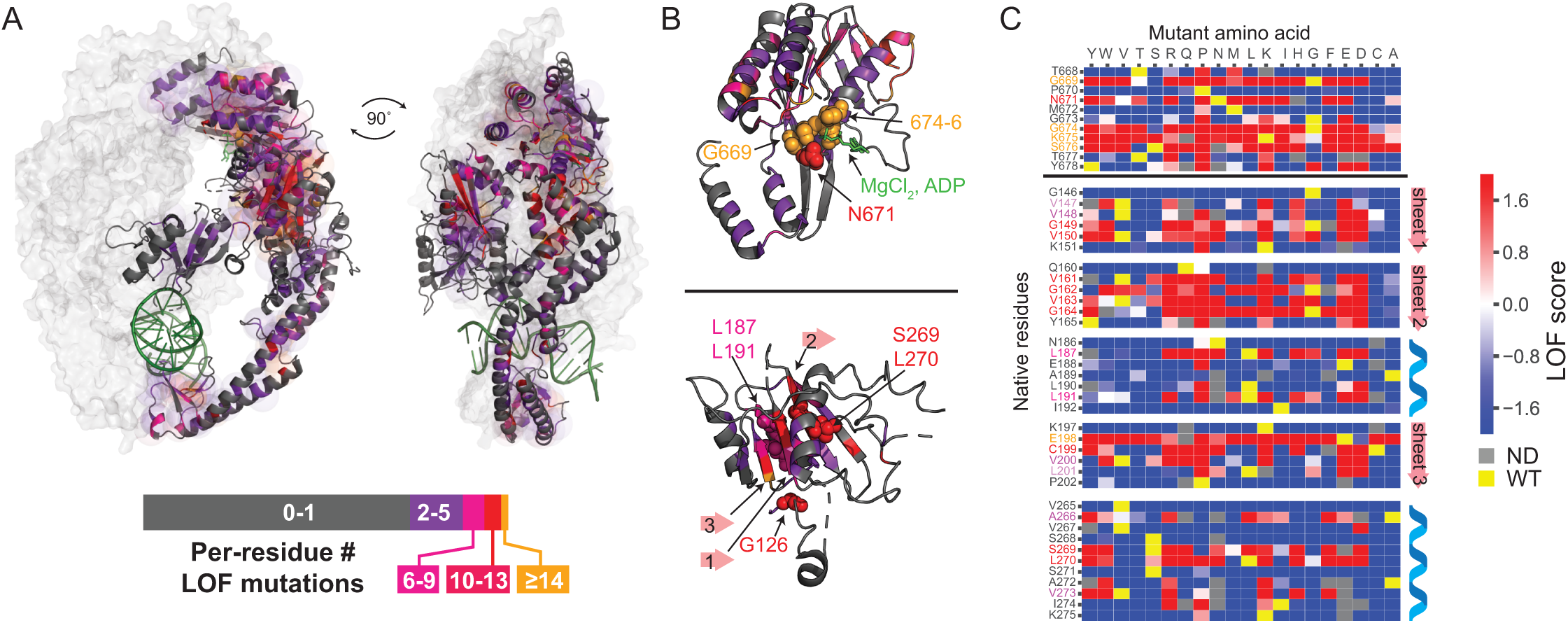
Structural insights from deep mutational scanning data. (A) MutSα crystal structure (PDB:2O8E), with MSH6 surface in light gray, and MSH2 shaded by the number of deleterious missense variants at each (length of bar in legend proportional to count of residues with denoted range of LOF mutations). (B) Extreme intolerance to substitution in MSH2 ATPase domain (upper), and connector domains (lower); selected residues or beta sheets shown with spheres or arrows, respectively. (C) LOF score heatmap (as in Figure 2C) showing mutational intolerance at Walker A motif of the ATPase domain (upper) and selected connector domain residues (lower) corresponding to (B).

We also examined the scores of variants accepted by evolution. Given the strong pressure to maintain genome integrity, *MSH2* variants rising to fixation in extant lineages are expected to be functionally neutral, except for potential epistatic effects of other mutations *in cis* or *trans*^*71*^. We aligned the human MSH2 protein sequence with seven orthologs distributed across eukaryota, identifying 1,707 amino acid substitutions relative to the human sequence. Of those scored, nearly all (1,627/1,639, 99.3%) had negative LOF scores (Figure S16), indicating these substitutions are functionally neutral in the context of the human MSH2 sequence, and supporting these scores’ specificity.

## Discussion

The mismatch repair factors underlying Lynch Syndrome are among the most intensively screened genes in clinical practice, and there is an acute need for functional evidence to assist in their variants’ interpretation as the scope of genetic testing broadens^72,73^. Massively parallel assays provide a tractable means to generate such evidence^74,75^. Here, we combined deep mutational scanning^17^ with an established MMR assay to generate a nearly complete functional effect map of missense variants in the key Lynch Syndrome gene *MSH2*.

The resulting LOF scores have high predictive accuracy, agreeing with traditional, small-scale functional reports for 158 of 165 previously tested missense variants (95.8%), which meets or exceeds the level of concordance among the previous functional screens of *MSH2* missense variants. To the best of our knowledge, MSH2 is the largest protein (934 aa) subjected to full-length deep mutational scanning to date, demonstrating that such pooled functional screens can scale to address the large space of clinically relevant human genetic variation. Additionally, this approach can be readily extended for comprehensive functional annotation of variants in other key Lynch Syndrome genes, *MLH1, PMS2*, and *MSH6*.

Our strategy exploited the direct link from molecular dysfunction - the inability to correct replicative errors - to pathogenesis in Lynch Syndrome. Accordingly, these functional measurements were highly consistent with expert-reviewed variant classification records from the ClinVar database and InSiGHT Group^9^. For two of the three conflicting variants, the standing classifications appear to be in error (Figure S13). The first, p.Glu198Gly (InSiGHT class 1: benign), was highly deleterious in our assay (LOF score: 3.47), a result corroborated by mutator assays in yeast^15,51^ and MNNG resistance following transient overexpression in human cells^49^. Additionally, all other substitutions at Glu198 appeared deleterious in our screen, except for the chemically similar p.Glu198Asp. The second example is p.Pro652His (InSiGHT class 4: likely pathogenic), which along with 14 of the other 15 measured substitutions at that codon, appeared neutral in our data, in agreement with its activity *in vitro*^*53*^. Thus, in addition to assisting the resolution of VUS’s, functional effect maps may allow identification and correction of rare misclassifications among clinical variation databases.

Overall, *MSH2* appeared quite tolerant to missense variation, with only 10.7% of assayed missense variants exhibiting loss-of-function. On face, this is surprising given the high level of sequence conservation among *MSH2* orthologs (e.g., ∼41% protein sequence identity between human and budding yeast), and given its conserved, essential role in mismatch repair. We consider it unlikely that large swaths of LOF variants have gone undetected given the strong concordance with expert-reviewed clinical interpretation and prior functional studies. This degree of tolerance is remarkable in the context of available deep mutational scanning data from other human genes. In terms of overall constraint, the dataset closest to *MSH2* is from the transcription factor *PPARG*, in which 21.5% of missense variants scored as having substantial probability of being casual for lipodystrophy^76^. Most other human genes subjected to full-length mutational scans thus far have shown substantially higher overall constraint: for instance, 40.5% of missense variants in the pharmacogene *NUDT15* scored as damaging^62^, as were 39% of variants in the homocysteine metabolic factor *CBS*^*77*^. A recent activity-agnostic protein stability screen showed the fraction of missense variants with substantially destabilizing effects to range from 25.1-43.1% across three different genes^62,78^; these could be taken as a lower bound given that an unknown fraction of true LOF variants may remain stable.

The bimodal distribution of LOF scores mirrors previous functional studies in which most *MSH2* missense variants have appeared either substantially functionally intact or null-like, with relatively few in between. An important question arising from these functional studies is whether variants with borderline scores reflect experimental noise or instead have intermediate functional activity^79^. For instance, several *MSH2* alleles modeled in yeast^15^ exhibited minor increases in reporter gene mutation rates (2-19X relative to *MSH2* WT, compared to >200X for *MSH2* null alleles), suggesting a mild MMR defect, but all of those scored as functionally neutral in our data. Of the three (p.L390F, p.M688I, and p.E886G) tested in other studies, all appeared functionally neutral, corroborating our results and suggesting either a greater sensitivity to functional defects in the yeast model or potentially some functional divergence between the orthologs^16^. Other variants have conflicting evidence which will require additional studies to resolve, for instance p.L93F, which showed intermediate activity (∼51%) in a cell-free MMR assay^53^, was undetected in a mouse ES cell based clonogenic screen for pathogenic alleles^19^ and was marginally neutral in our data (LOF score: -0.26).

For any such alleles that do behave as hypomorphs in functional assays, it remains unclear whether these meaningfully contribute to Lynch Syndrome risk^51,80^. Complicating this question is the challenge of establishing a truth set of moderate-risk *MSH2* variants in family or case-control studies, as with further reduced penetrance and later age of cancer onset, these may resemble common sporadic cancers. However, anecdotes suggest that even severely hypomorphic alleles of MMR factors may retain enough activity to set them apart from complete LOF alleles. For instance, a splice-disruptive mutation which reduces PMS2 expression by >10-fold retains enough activity that when homozygous, it results in a form of constitutional MMR deficiency which is highly attenuated, with later onset and less profound microsatellite instability compared to truly null biallelic mutations^81^. As increasingly large variation datasets coupled to electronic health records become available^82–84^, it may become possible to examine whether variants with intermediate functional scores, on aggregate, confer increased Lynch Syndrome risk, and whether individuals carrying one of these alleles opposite a *bona fide* pathogenic allele present with biallelic MMR deficiency.

One limitation of these cDNA-based measurements is that they cannot model effects upon splicing, although among known Lynch Syndrome variants and in saturation screens of other hereditary cancer syndrome genes^85^, splice disruption accounts for only a small fraction of pathogenic variants. In the future, splicing effects could be accounted for by incorporating improved bioinformatic predictions^31^ or multiplexed assays^86–88^. Another potential limitation is that our *MSH2* LOF scores are not a direct measure of mutation rates associated with each variant, to the extent that these could differ across *MSH2* variants. We instead used a close proxy for MMR dysfunction, treatment with the purine analog 6-TG, which selectively enrichies for MMR-defective cells which lack the ability to remove the resulting lesions. As with other deep mutational scans, these results may be further refined with orthogonal functional selections^62,89^ or complemented by specialized assays for protein stability^78^, localization^90^, or partner (MSH6) binding.

We demonstrate that massively parallel functional assays can accurately measure the impacts of variants in the key Lynch Syndrome gene *MSH2*. This functional effect map, if rationally combined^54,91^ with other lines of clinical evidence^9^, will enable more accurate classification of Lynch Syndrome variants. Adding confidence and accuracy to Lynch Syndrome variant interpretation will afford carriers of pathogenic variants the benefits of potentially life-saving early detection and medical interventions^92^.

## Supplemental Data

Supplemental Data include sixteen figures and five tables

## Acknowledgements

We thank M. Meisler, A. Antonellis, E. Stoffel, S. Hemker, and A. Glazer for helpful discussions, J. Moran for providing sequencer access, and J. Moldovan for assistance with clonogenic assays. This work was supported by the National Institute of General Medical Sciences (5R01GM129123 to J.O.K.) and a Precision Health Fellowship from the University of Michigan (to X.J.).

## Declaration of Interests

The authors declare no competing interests.

## Notes

### Competing Interest Statement

The authors have declared no competing interest.

## References

1. Lynch, H.T., and Krush, A.J. (1971). Cancer family “G” revisited: 1895-1970. Cancer 27, 1505–1511.

2. Leach, F.S., Nicolaides, N.C., Papadopoulos, N., Liu, B., Jen, J., Parsons, R., Peltomäki, P., Sistonen, P., Aaltonen, L.A., and Nyström-Lahti, M. (1993). Mutations of a mutS homolog in hereditary nonpolyposis colorectal cancer. Cell 75, 1215–1225.

3. Fishel, R., Lescoe, M.K., Rao, M.R., Copeland, N.G., Jenkins, N.A., Garber, J., Kane, M., and Kolodner, R. (1994). The human mutator gene homolog MSH2 and its association with hereditary nonpolyposis colon cancer. Cell 77, 1 p following 166.

4. Bronner, C.E., Baker, S.M., Morrison, P.T., Warren, G., Smith, L.G., Lescoe, M.K., Kane, M., Earabino, C., Lipford, J., Lindblom, A., et al. (1994). Mutation in the DNA mismatch repair gene homologue hMLH 1 is associated with hereditary non-polyposis colon cancer. Nature 368, 258–261.

5. Papadopoulos, N., Nicolaides, N.C., Wei, Y.F., Ruben, S.M., Carter, K.C., Rosen, C.A., Haseltine, W.A., Fleischmann, R.D., Fraser, C.M., and Adams, M.D. (1994). Mutation of a mutL homolog in hereditary colon cancer. Science 263, 1625–1629.

6. Win, A.K., Jenkins, M.A., Dowty, J.G., Antoniou, A.C., Lee, A., Giles, G.G., Buchanan, D.D., Clendenning, M., Rosty, C., Ahnen, D.J., et al. (2017). Prevalence and Penetrance of Major Genes and Polygenes for Colorectal Cancer. Cancer Epidemiol. Biomarkers Prev. 26, 404–412.

7. Haraldsdottir, S., Rafnar, T., Frankel, W.L., Einarsdottir, S., Sigurdsson, A., Hampel, H., Snaebjornsson, P., Masson, G., Weng, D., Arngrimsson, R., et al. (2017). Comprehensive population-wide analysis of Lynch syndrome in Iceland reveals founder mutations in MSH6 and PMS2. Nat. Commun. 8, 14755.

8. Kalia, S.S., Adelman, K., Bale, S.J., Chung, W.K., Eng, C., Evans, J.P., Herman, G.E., Hufnagel, S.B., Klein, T.E., Korf, B.R., et al. (2017). Recommendations for reporting of secondary findings in clinical exome and genome sequencing, 2016 update (ACMG SF v2. 0): a policy statement of the American College of Medical Genetics and Genomics. Genet. Med. 19, 249.

9. Thompson, B.A., Spurdle, A.B., Plazzer, J.-P., Greenblatt, M.S., Akagi, K., Al-Mulla, F., Bapat, B., Bernstein, I., Capellá, G., den Dunnen, J.T., et al. (2014). Application of a 5-tiered scheme for standardized classification of 2,360 unique mismatch repair gene variants in the InSiGHT locus-specific database. Nat. Genet. 46, 107–115.

10. Giardiello, F.M., Allen, J.I., Axilbund, J.E., Boland, C.R., Burke, C.A., Burt, R.W., Church, J.M., Dominitz, J.A., Johnson, D.A., Kaltenbach, T., et al. (2014). Guidelines on genetic evaluation and management of Lynch syndrome: a consensus statement by the US Multi-Society Task Force on Colorectal Cancer. CDis. olon Rectum 57, 1025–1048.

11. Møller, P., Seppälä, T.T., Bernstein, I., Holinski-Feder, E., Sala, P., Gareth Evans, D., Lindblom, A., Macrae, F., Blanco, I., Sijmons, R.H., et al. (2018). Cancer risk and survival in path_MMR carriers by gene and gender up to 75 years of age: a report from the Prospective Lynch Syndrome Database. Gut 67, 1306–1316.

12. Brnich, S.E., Rivera-Muñoz, E.A., and Berg, J.S. (2018). Quantifying the potential of functional evidence to reclassify variants of uncertain significance in the categorical and Bayesian interpretation frameworks. CHum. Mutat. 39, 1531–1541.

13. Richards, S., Aziz, N., Bale, S., Bick, D., Das, S., Gastier-Foster, J., Grody, W.W., Hegde, M., Lyon, E., Spector, E., et al. (2015). Standards and guidelines for the interpretation of sequence variants: a joint consensus recommendation of the American College of Medical Genetics and Genomics and the Association for Molecular Pathology. CGenet. Med. 17, 405–424.

14. Clark, A.B., Cook, M.E., Tran, H.T., Gordenin, D.A., Resnick, M.A., and Kunkel, T.A. (1999). Functional analysis of human MutSα and MutSβ complexes in yeast. CNucleic Acids Res. 27, 736–742.

15. Gammie, A.E., Erdeniz, N., Beaver, J., Devlin, B., Nanji, A., and Rose, M.D. (2007). Functional characterization of pathogenic human MSH2 missense mutations in Saccharomyces cerevisiae. Genetics 177, 707–721.

16. Shimodaira, H., Filosi, N., Shibata, H., Suzuki, T., Radice, P., Kanamaru, R., Friend, S.H., Kolodner, R.D., and Ishioka, C. (1998). Functional analysis of human MLH1 mutations in Saccharomyces cerevisiae. CNat. Genet. 19, 384–389.

17. Fowler, D.M., and Fields, S. (2014). Deep mutational scanning: a new style of protein science. CNat. Methods 11, 801–807.

18. Swann, P.F., Waters, T.R., Moulton, D.C., Xu, Y.Z., Zheng, Q., Edwards, M., and Mace, R. (1996). Role of postreplicative DNA mismatch repair in the cytotoxic action of thioguanine. Science 273, 1109–1111.

19. Houlleberghs, H., Dekker, M., Lantermans, H., Kleinendorst, R., Dubbink, H.J., Hofstra, R.M.W., Verhoef, S., and Te Riele, H. (2016). Oligonucleotide-directed mutagenesis screen to identify pathogenic Lynch syndrome-associated MSH2 DNA mismatch repair gene variants. Proc. Natl. Acad. Sci. U. S. A. 113, 4128–4133.

20. Yan, T., Berry, S.E., Desai, A.B., and Kinsella, T.J. (2003). DNA mismatch repair (MMR) mediates 6-thioguanine genotoxicity by introducing single-strand breaks to signal a G2-M arrest in MMR-proficient RKO cells. Clin. Cancer Res. 9, 2327–2334.

21. Mojas, N., Lopes, M., and Jiricny, J. (2007). Mismatch repair-dependent processing of methylation damage gives rise to persistent single-stranded gaps in newly replicated DNA. Genes Dev. 21, 3342–3355.

22. Aguirre, M., Rivas, M.A., and Priest, J. (2019). Phenome-wide Burden of Copy-Number Variation in the UK Biobank. Am. J. Hum. Genet. 105, 373–383.

23. Karczewski, K.J., Francioli, L.C., Tiao, G., Cummings, B.B., Alföldi, J., Wang, Q., Collins, R.L., Laricchia, K.M., Ganna, A., Birnbaum, D.P., et al. (2020). The mutational constraint spectrum quantified from variation in 141,456 humans. Nature 581, 434–443.

24. Ran, F.A., Hsu, P.D., Wright, J., Agarwala, V., Scott, D.A., and Zhang, F. (2013). Genome engineering using the CRISPR-Cas9 system. Nat. Protoc. 8, 2281–2308.

25. Kitzman, J.O., Starita, L.M., Lo, R.S., Fields, S., and Shendure, J. (2015). Massively parallel single-amino-acid mutagenesis. Nat. Methods 12, 203–206, 4 p following 206.

26. Rohland, N., and Reich, D. (2012). Cost-effective, high-throughput DNA sequencing libraries for multiplexed target capture. Genome Res. 22, 939–946.

27. Picelli, S., Björklund, A.K., Reinius, B., Sagasser, S., Winberg, G., and Sandberg, R. (2014). Tn5 transposase and tagmentation procedures for massively scaled sequencing projects. Genome Res. 24, 2033–2040.

28. Joung, J., Konermann, S., Gootenberg, J.S., Abudayyeh, O.O., Platt, R.J., Brigham, M.D., Sanjana, N.E., and Zhang, F. (2017). Genome-scale CRISPR-Cas9 knockout and transcriptional activation screening. Nat. Protoc. 12, 828–863.

29. Zhang, J., Kobert, K., Flouri, T., and Stamatakis, A. (2014). PEAR: a fast and accurate Illumina Paired-End reAd mergeR. Bioinformatics 30, 614–620.

30. Zorita, E., Cuscó, P., and Filion, G.J. (2015). Starcode: sequence clustering based on all-pairs search. Bioinformatics 31, 1913–1919.

31. Jaganathan, K., Kyriazopoulou Panagiotopoulou, S., McRae, J.F., Darbandi, S.F., Knowles, D., Li, Y.I., Kosmicki, J.A., Arbelaez, J., Cui, W., Schwartz, G.B., et al. (2019). Predicting Splicing from Primary Sequence with Deep Learning. Cell 176, 535–548.e24.

32. Carlson, J., Locke, A.E., Flickinger, M., Zawistowski, M., Levy, S., Myers, R.M., Boehnke, M., Kang, H.M., Scott, L.J., Li, J.Z., et al. (2018). Extremely rare variants reveal patterns of germline mutation rate heterogeneity in humans. Nat. Commun. 9, 3753.

33. Rentzsch, P., Witten, D., Cooper, G.M., Shendure, J., and Kircher, M. (2019). CADD: predicting the deleteriousness of variants throughout the human genome. Nucleic Acids Res. 47, D886–D894.

34. Adzhubei, I.A., Schmidt, S., Peshkin, L., Ramensky, V.E., Gerasimova, A., Bork, P., Kondrashov, A.S., and Sunyaev, S.R. (2010). A method and server for predicting damaging missense mutations. Nat. Methods 7, 248–249.

35. Ioannidis, N.M., Rothstein, J.H., Pejaver, V., Middha, S., McDonnell, S.K., Baheti, S., Musolf, A., Li, Q., Holzinger, E., Karyadi, D., et al. (2016). REVEL: An Ensemble Method for Predicting the Pathogenicity of Rare Missense Variants. Am. J. Hum. Genet. 99, 877–885.

36. Ali, H., Olatubosun, A., and Vihinen, M. (2012). Classification of mismatch repair gene missense variants with PON-MMR. Hum. Mutat. 33, 642–650.

37. Chao, E., Velasquez, J., Witherspoon, M., Rozek, L., Ng, P., Gruber, S., Watson, P., Rennert, G., Anton-Culver, H., Lynch, H., et al. (2008). Accurate classification of missense variants in MLH1/MSH2 with MAPP-MMR. Cancer Res. 68, LB – 14 – LB – 14.

38. Nielsen, S.V., Stein, A., Dinitzen, A.B., Papaleo, E., Tatham, M.H., Poulsen, E.G., Kassem, M.M., Rasmussen, L.J., Lindorff-Larsen, K., and Hartmann-Petersen, R. (2017). Predicting the impact of Lynch syndrome-causing missense mutations from structural calculations. PLoS Genet. 13, e1006739.

39. Warren, J.J., Pohlhaus, T.J., Changela, A., Iyer, R.R., Modrich, P.L., and Beese, L.S. (2007). Structure of the Human MutSα DNA Lesion Recognition Complex. Mol. Cell 26, 579–592.

40. Touw, W.G., Baakman, C., Black, J., te Beek, T.A.H., Krieger, E., Joosten, R.P., and Vriend, G. (2015). A series of PDB-related databanks for everyday needs. Nucleic Acids Research 43, D364–D368.

41. Edgar, R.C. (2004). MUSCLE: multiple sequence alignment with high accuracy and high throughput. Nucleic Acids Res. 32, 1792–1797.

42. Harmsen, T., Klaasen, S., van de Vrugt, H., and Te Riele, H. (2018). DNA mismatch repair and oligonucleotide end-protection promote base-pair substitution distal from a CRISPR/Cas9-induced DNA break. Nucleic Acids Res. 46, 2945–2955.

43. Foulkes, W.D., Thiffault, I., Gruber, S.B., Horwitz, M., Hamel, N., Lee, C., Shia, J., Markowitz, A., Figer, A., Friedman, E., et al. (2002). The Founder Mutation MSH2*1906G?C Is an Important Cause of Hereditary Nonpolyposis Colorectal Cancer in the Ashkenazi Jewish Population. Am. J. Hum. Genet. 71, 1395–1412.

44. Wielders, E.A.L., Hettinger, J., Dekker, R., Kets, C.M., Ligtenberg, M.J., Mensenkamp, A.R., van den Ouweland, A.M.W., Prins, J., Wagner, A., Dinjens, W.N.M., et al. (2014). Functional analysis of MSH2 unclassified variants found in suspected Lynch syndrome patients reveals pathogenicity due to attenuated mismatch repair. J. Med. Genet. 51, 245–253.

45. Weile, J., Sun, S., Cote, A.G., Knapp, J., Verby, M., Mellor, J.C., Wu, Y., Pons, C., Wong, C., van Lieshout, N., et al. (2017). A framework for exhaustively mapping functional missense variants. Mol. Syst. Biol. 13, 957.

46. Sack, L.M., Davoli, T., Xu, Q., Li, M.Z., and Elledge, S.J. (2016). Sources of Error in Mammalian Genetic Screens. G3 6, 2781–2790.

47. Rath, A., Mishra, A., Ferreira, V.D., Hu, C., Omerza, G., Kelly, K., Hesse, A., Reddi, H.V., Grady, J.P., and Heinen, C.D. (2019). Functional interrogation of Lynch syndrome-associated MSH2 missense variants via CRISPR-Cas9 gene editing in human embryonic stem cells. Hum. Mutat. 40, 2044–2056.

48. Mastrocola, A.S., and Heinen, C.D. (2010). Lynch syndrome-associated mutations in MSH2 alter DNA repair and checkpoint response functions in vivo. Hum. Mutat. 31, E1699–E1708.

49. Bouvet, D., Bodo, S., Munier, A., Guillerm, E., Bertrand, R., Colas, C., Duval, A., Coulet, F., and Muleris, M. (2019). Methylation Tolerance-Based Functional Assay to Assess Variants of Unknown Significance in the MLH1 and MSH2 Genes and Identify Patients With Lynch Syndrome. Gastroenterology 157, 421–431.

50. Drost, M., Lützen, A., van Hees, S., Ferreira, D., Calléja, F., Zonneveld, J.B.M., Nielsen, F.C., Rasmussen, L.J., and de Wind, N. (2013). Genetic screens to identify pathogenic gene variants in the common cancer predisposition Lynch syndrome. Proc. Natl. Acad. Sci. U. S. A. 110, 9403–9408.

51. Martinez, S.L., and Kolodner, R.D. (2010). Functional analysis of human mismatch repair gene mutations identifies weak alleles and polymorphisms capable of polygenic interactions. Proc. Natl. Acad. Sci. U. S. A. 107, 5070–5075.

52. Drost, M., Zonneveld, J.B.M., van Hees, S., Rasmussen, L.J., Hofstra, R.M.W., and de Wind, N. (2012). A rapid and cell-free assay to test the activity of lynch syndrome-associated MSH2 and MSH6 missense variants. Human Mutation 33, 488–494.

53. Drost, M., Tiersma, Y., Thompson, B.A., Frederiksen, J.H., Keijzers, G., Glubb, D., Kathe, S., Osinga, J., Westers, H., Pappas, L., et al. (2019). A functional assay–based procedure to classify mismatch repair gene variants in Lynch syndrome. Genet. Med. 21, 1486–1496.

54. Brnich, S.E., Abou Tayoun, A.N., Couch, F.J., Cutting, G.R., Greenblatt, M.S., Heinen, C.D., Kanavy, D.M., Luo, X., McNulty, S.M., Starita, L.M., et al. (2019). Recommendations for application of the functional evidence PS3/BS3 criterion using the ACMG/AMP sequence variant interpretation framework. Genome Med. 12, 3.

55. Chao, E.C., Velasquez, J.L., Witherspoon, M.S.L., Rozek, L.S., Peel, D., Ng, P., Gruber, S.B., Watson, P., Rennert, G., Anton-Culver, H., et al. (2008). Accurate classification of MLH1/MSH2 missense variants with multivariate analysis of protein polymorphisms-mismatch repair (MAPP-MMR). Hum. Mutat. 29, 852–860.

56. Schymkowitz, J., Borg, J., Stricher, F., Nys, R., Rousseau, F., and Serrano, L. (2005). The FoldX web server: an online force field. Nucleic Acids Res. 33, W382–W388.

57. Ali, H., Olatubosun, A., and Vihinen, M. (2012). Classification of mismatch repair gene missense variants with PON-MMR. Hum. Mutat. 33, 642–650.

58. Kircher, M., Witten, D.M., Jain, P., O’Roak, B.J., Cooper, G.M., and Shendure, J. (2014). A general framework for estimating the relative pathogenicity of human genetic variants. Nat. Genet. 46, 310–315.

59. Chicco, D., and Jurman, G. (2020). The advantages of the Matthews correlation coefficient (MCC) over F1 score and accuracy in binary classification evaluation. BMC Genomics 21, 6.

60. van der Velde, K.J., Kuiper, J., Thompson, B.A., Plazzer, J.-P., van Valkenhoef, G., de Haan, M., Jongbloed, J.D.H., Wijmenga, C., de Koning, T.J., Abbott, K.M., et al. (2015). Evaluation of CADD Scores in Curated Mismatch Repair Gene Variants Yields a Model for Clinical Validation and Prioritization. Hum. Mutat. 36, 712–719.

61. Li, J., Zhao, T., Zhang, Y., Zhang, K., Shi, L., Chen, Y., Wang, X., and Sun, Z. (2018). Performance evaluation of pathogenicity-computation methods for missense variants. Nucleic Acids Res. 46, 7793–7804.

62. Suiter, C.C., Moriyama, T., Matreyek, K.A., Yang, W., Scaletti, E.R., Nishii, R., Yang, W., Hoshitsuki, K., Singh, M., Trehan, A., et al. (2020). Massively parallel variant characterization identifies NUDT15 alleles associated with thiopurine toxicity. Proc. Natl. Acad. Sci. U. S. A.

63. Livesey, B.J., and Marsh, J.A. (2020). Using deep mutational scanning to benchmark variant effect predictors and identify disease mutations. Mol. Syst. Biol. 16, e9380.

64. Gray, V.E., Hause, R.J., and Fowler, D.M. (2017). Analysis of Large-Scale Mutagenesis Data To Assess the Impact of Single Amino Acid Substitutions. Genetics 207, 53–61.

65. Rollins, N.J., Brock, K.P., Poelwijk, F.J., Stiffler, M.A., Gauthier, N.P., Sander, C., and Marks, D.S. (2019). Inferring protein 3D structure from deep mutation scans. Nat. Genet. 51, 1170–1176.

66. Graham, W.J., Putnam, C.D., and Kolodner, R.D. (2018). The properties of Msh2–Msh6 ATP binding mutants suggest a signal amplification mechanism in DNA mismatch repair. J. Biol. Chem.

67. Kumar, C., Piacente, S.C., Sibert, J., Bukata, A.R., O’Connor, J., Alani, E., and Surtees, J.A. (2011). Multiple Factors Insulate Msh2–Msh6 Mismatch Repair Activity from Defects in Msh2 Domain I. J. Mol. Biol. 411, 765–780.

68. Cyr, J.L., Brown, G.D., Stroop, J., and Heinen, C.D. (2012). The predicted truncation from a cancer-associated variant of the MSH2 initiation codon alters activity of the MSH2-MSH6 mismatch repair complex. Molecular Carcinogenesis 51, 647–658.

69. Kets, C.M., Hoogerbrugge, N., van Krieken, J.H.J.M., Goossens, M., Brunner, H.G., and Ligtenberg, M.J.L. (2009). Compound heterozygosity for two MSH2 mutations suggests mild consequences of the initiation codon variant c.1A>G of MSH2. Eur. J. Hum. Genet. 17, 159–164.

70. Obmolova, G., Ban, C., Hsieh, P., and Yang, W. (2000). Crystal structures of mismatch repair protein MutS and its complex with a substrate DNA. Nature 407, 703–710.

71. Jordan, D.M., Frangakis, S.G., Golzio, C., Cassa, C.A., Kurtzberg, J., Task Force for Neonatal Genomics, Davis, E.E., Sunyaev, S.R., and Katsanis, N. (2015). Identification of cis-suppression of human disease mutations by comparative genomics. Nature 524, 225–229.

72. Dinh, T.A., Rosner, B.I., Atwood, J.C., Boland, C.R., Syngal, S., Vasen, H.F.A., Gruber, S.B., and Burt, R.W. (2011). Health benefits and cost-effectiveness of primary genetic screening for Lynch syndrome in the general population. Cancer Prev. Res. 4, 9–22.

73. King, M.-C., Levy-Lahad, E., and Lahad, A. (2014). Population-based screening for BRCA1 and BRCA2: 2014 Lasker Award. JAMA 312, 1091–1092.

74. Starita, L.M., Ahituv, N., Dunham, M.J., Kitzman, J.O., Roth, F.P., Seelig, G., Shendure, J., and Fowler, D.M. (2017). Variant Interpretation: Functional Assays to the Rescue. Am. J. Hum. Genet. 101, 315–325.

75. Weile, J., and Roth, F.P. (2018). Multiplexed assays of variant effects contribute to a growing genotype-phenotype atlas. Hum. Genet. 137, 665–678.

76. Majithia, A.R., Tsuda, B., Agostini, M., Gnanapradeepan, K., Rice, R., Peloso, G., Patel, K.A., Zhang, X., Broekema, M.F., Patterson, N., et al. (2016). Prospective functional classification of all possible missense variants in PPARG. Nat. Genet. 48, 1570–1575.

77. Sun, S., Weile, J., Verby, M., Wu, Y., Wang, Y., Cote, A.G., Fotiadou, I., Kitaygorodsky, J., Vidal, M., Rine, J., et al. (2020). A proactive genotype-to-patient-phenotype map for cystathionine beta-synthase. Genome Med. 12, 13.

78. Matreyek, K.A., Starita, L.M., Stephany, J.J., Martin, B., Chiasson, M.A., Gray, V.E., Kircher, M., Khechaduri, A., Dines, J.N., Hause, R.J., et al. (2018). Multiplex assessment of protein variant abundance by massively parallel sequencing. Nat. Genet. 50, 874–882.

79. Spurdle, A.B., Whiley, P.J., Thompson, B., Feng, B., Healey, S., Brown, M.A., Pettigrew, C., kConFab, Van Asperen, C.J., Ausems, M.G.E.M., et al. (2012). BRCA1 R1699Q variant displaying ambiguous functional abrogation confers intermediate breast and ovarian cancer risk. J. Med. Genet. 49, 525–532.

80. Heinen, C.D., and Juel Rasmussen, L. (2012). Determining the functional significance of mismatch repair gene missense variants using biochemical and cellular assays. Hered. Cancer Clin. Pract. 10, 9.

81. Li, L., Hamel, N., Baker, K., McGuffin, M.J., Couillard, M., Gologan, A., Marcus, V.A., Chodirker, B., Chudley, A., Stefanovici, C., et al. (2015). A homozygous PMS2 founder mutation with an attenuated constitutional mismatch repair deficiency phenotype. J. Med. Genet. 52, 348–352.

82. Carey, D.J., Fetterolf, S.N., Davis, F.D., Faucett, W.A., Kirchner, H.L., Mirshahi, U., Murray, M.F., Smelser, D.T., Gerhard, G.S., and Ledbetter, D.H. (2016). The Geisinger MyCode community health initiative: an electronic health record–linked biobank for precision medicine research. Genet. Med. 18, 906–913.

83. Bycroft, C., Freeman, C., Petkova, D., Band, G., Elliott, L.T., Sharp, K., Motyer, A., Vukcevic, D., Delaneau, O., O’Connell, J., et al. (2018). The UK Biobank resource with deep phenotyping and genomic data. Nature 562, 203–209.

84. Grzymski, J.J., Elhanan, G., Morales Rosado, J.A., Smith, E., Schlauch, K.A., Read, R., Rowan, C., Slotnick, N., Dabe, S., Metcalf, W.J., et al. (2020). Population genetic screening efficiently identifies carriers of autosomal dominant diseases. Nat. Med.

85. Findlay, G.M., Daza, R.M., Martin, B., Zhang, M.D., Leith, A.P., Gasperini, M., Janizek, J.D., Huang, X., Starita, L.M., and Shendure, J. (2018). Accurate classification of BRCA1 variants with saturation genome editing. Nature 562, 217–222.

86. Adamson, S.I., Zhan, L., and Graveley, B.R. (2018). Vex-seq: high-throughput identification of the impact of genetic variation on pre-mRNA splicing efficiency. Genome Biol. 19, 71.

87. Cheung, R., Insigne, K.D., Yao, D., Burghard, C.P., Wang, J., Hsiao, Y.-H.E., Jones, E.M., Goodman, D.B., Xiao, X., and Kosuri, S. (2019). A Multiplexed Assay for Exon Recognition Reveals that an Unappreciated Fraction of Rare Genetic Variants Cause Large-Effect Splicing Disruptions. Mol. Cell 73, 183–194.e8.

88. Soemedi, R., Cygan, K.J., Rhine, C.L., Wang, J., Bulacan, C., Yang, J., Bayrak-Toydemir, P., McDonald, J., and Fairbrother, W.G. (2017). Pathogenic variants that alter protein code often disrupt splicing. Nat. Genet. 49, 848–855.

89. Mighell, T.L., Thacker, S., Fombonne, E., Eng, C., and O’Roak, B.J. (2020). An Integrated Deep-Mutational-Scanning Approach Provides Clinical Insights on PTEN Genotype-Phenotype Relationships. Am. J. Hum. Genet. 106, 818–829.

90. Penn, W.D., McKee, A.G., Kuntz, C.P., Woods, H., Nash, V., Gruenhagen, T.C., Roushar, F.J., Chandak, M., Hemmerich, C., Rusch, D.B., et al. (2020). Probing biophysical sequence constraints within the transmembrane domains of rhodopsin by deep mutational scanning. Sci Adv 6, eaay7505.

91. Tavtigian, S.V., Greenblatt, M.S., Harrison, S.M., Nussbaum, R.L., Prabhu, S.A., Boucher, K.M., Biesecker, L.G., and ClinGen Sequence Variant Interpretation Working Group (ClinGen SVI) (2018). Modeling the ACMG/AMP variant classification guidelines as a Bayesian classification framework. Genet. Med. 20, 1054–1060.

92. Hampel, H., and de la Chapelle, A. (2011). The search for unaffected individuals with Lynch syndrome: do the ends justify the means? Cancer Prev. Res. 4, 1–5.

