## Supplementary Figures for "Massively parallel functional testing of *MSH2* missense variants conferring Lynch Syndrome risk"

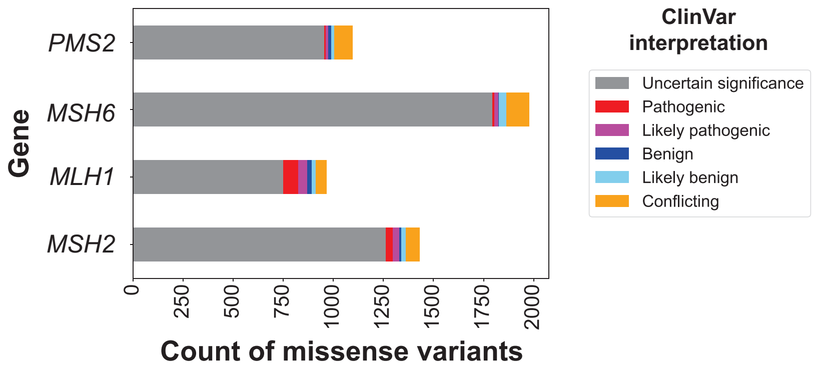


**Figure S1. Clinical missense variants in Lynch Syndrome genes.**
Counts of missense variants listed in the ClinVar database are shown by gene and shaded by classification.

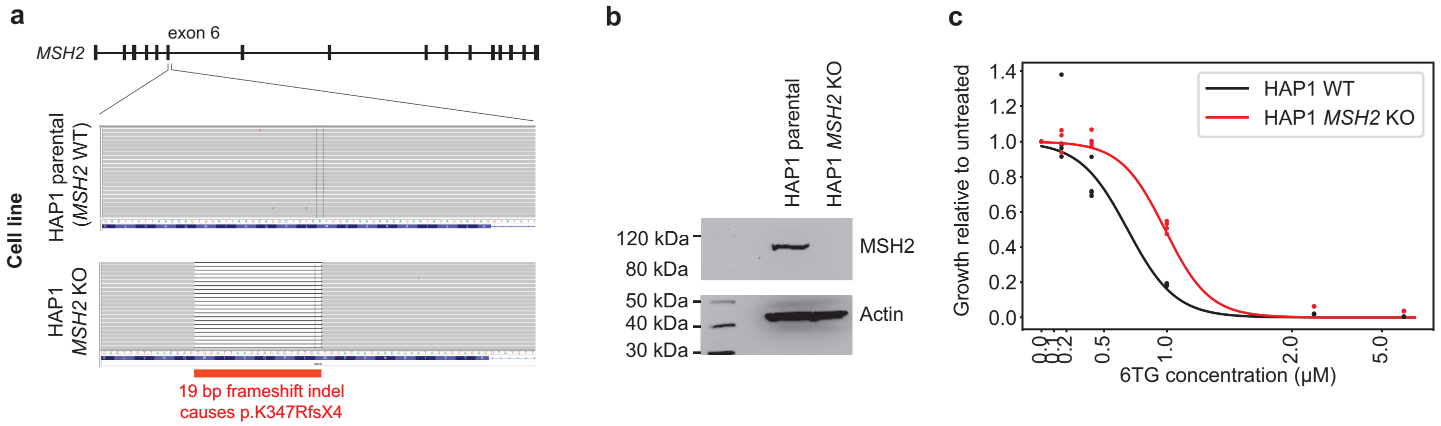


**Figure S2. HAP1 *MSH2* knockout cell line.**

**a**, Genotyping of *MSH2* exon 6 in HAP1 wild type cells (upper) and knockout cells (lower), showing a 19-bp frameshifting deletion introduced by CRISPR-Cas9 genome editing in the latter. **b**, Western blot of MSH2 (actin loading control) in parental and *MSH2* KO HAP1 lines. **c**, Dose response curve showing proliferation following 96 hours 6-TG treatment, relative to no treatment for KO (red) and WT (black) cells.

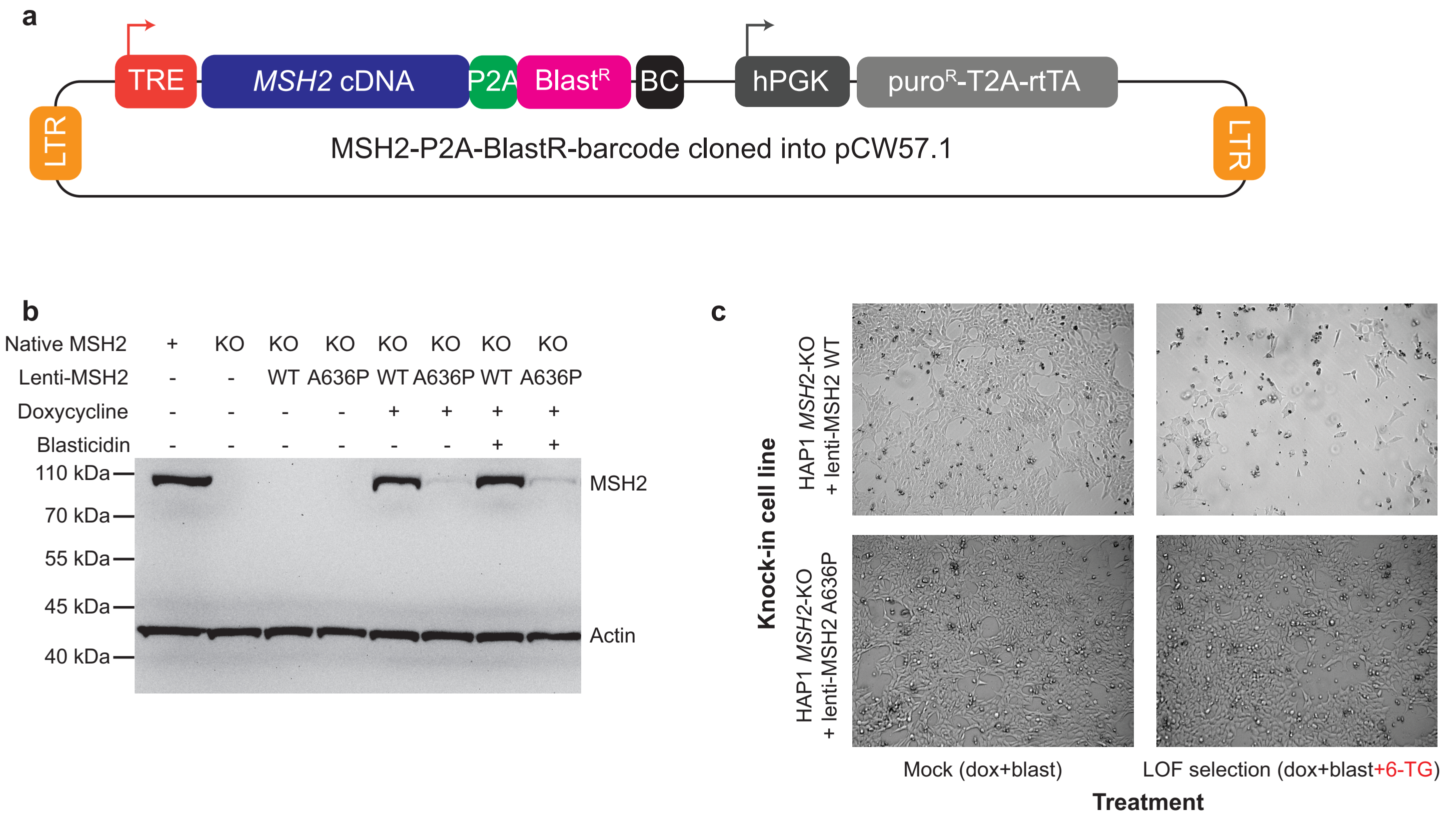


**Figure S3. Stable transduction with *MSH2* cDNA and functional selection with 6-TG.**
**a**, Schematic of doxycycline-inducible 'Tet-On' lentiviral transfer plasmid used to stably integrate *MSH2* cDNA and barcode (BC) into *MSH2* KO cells. **b**, Western blot of MSH2 (and actin loading control) under basal conditions and after induction with doxycycline. Upon induction, MSH2 A636P is only faintly detected compared with WT MSH2, but both gain blasticidin resistance (rightmost two columns). **c**, Representative images of *MSH2* knock-in cell lines, HAP1 *MSH2* KO+WT (top row) and HAP1 *MSH2* KO+A636P (bottom row), following mock treatment (left column: doxycycline, blasticidin, puromycin) or 6-TG selective conditions (right column: doxycycline, blasticidin, puromycin, 6-TG).

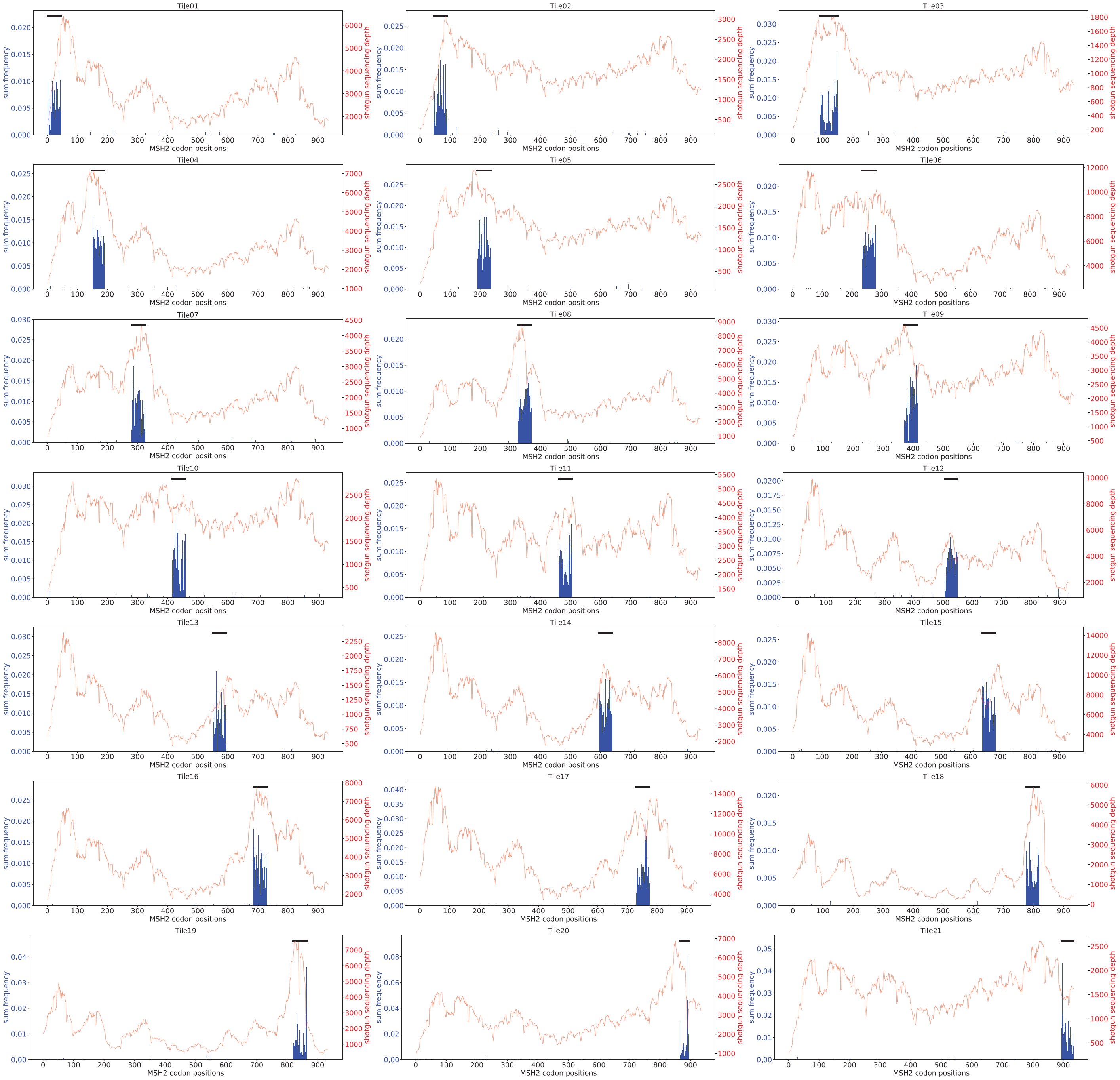


**Figure S4. Specificity of mutagenesis**.

Per-codon mutation rate was estimated by deep shotgun sequencing of all 21 integrated *MSH2* mutant tile libraries. Plotted are aggregated mutation rates (blue) for each codon position with edit distance 2 or 3 mutations, and sequencing depth (red). Black bars indicate tile positions.

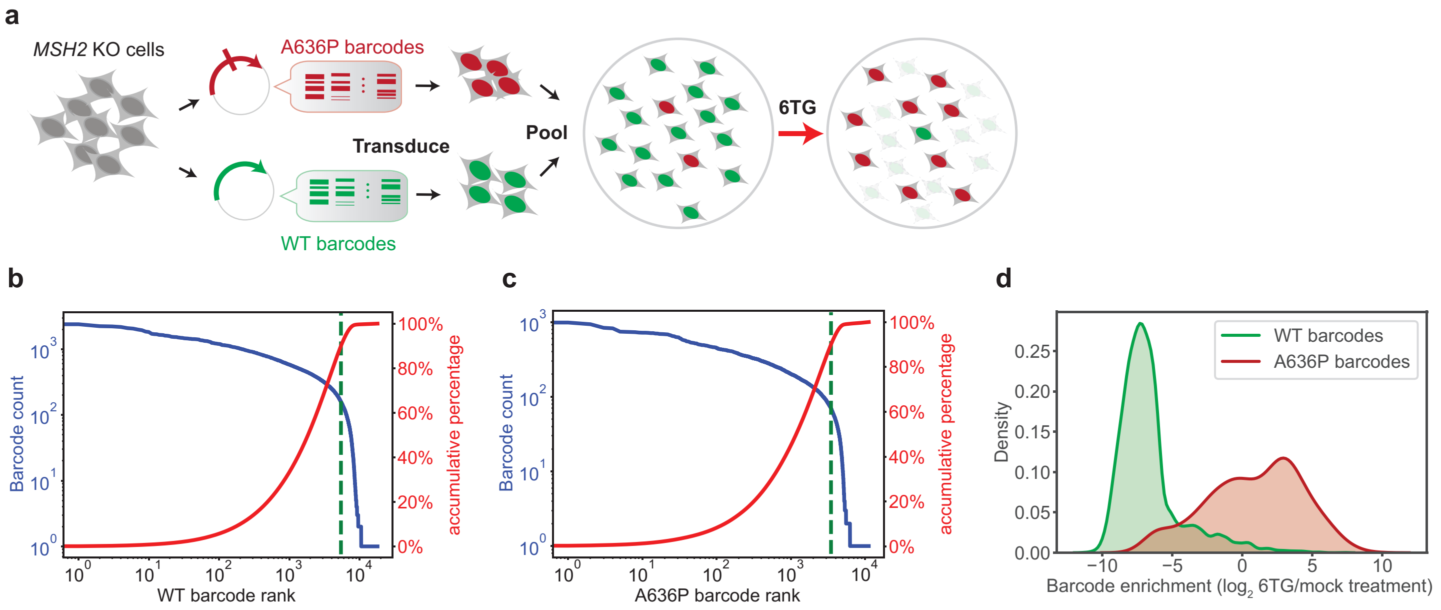


**Figure S5. Pooled functional selection using known *MSH2* variants.**
**a**, HAP1 *MSH2* KO cells were separately transduced with *MSH2* cDNA lentivirus (WT: green, A636P: red), each bearing libraries of barcodes. Resulting WT and A636P knock-in cells were mixed, then selected with 6-TG, and integrated barcodes were sequenced. **b**, **c**, Waterfall plot of integrated barcode abundances before treatment, showing barcode frequency versus rank (blue curve; left y-axis) and cumulative frequency vs rank (red curve; right y-axis), in *MSH2* WT (**b**) and A636P (**c**) knock-in cells. Dotted line denotes 90th percentile (i.e., rank of barcode at which all barcodes with greater or equal abundance comprise 90% of counts). No overlap was observed between WT and A636P barcode populations. **d**, Density plot of individual barcodes' log2 fold change after 6-TG treatment relative to mock, for barcodes associated with WT *MSH2* (green) and A636P *MSH2* (red).

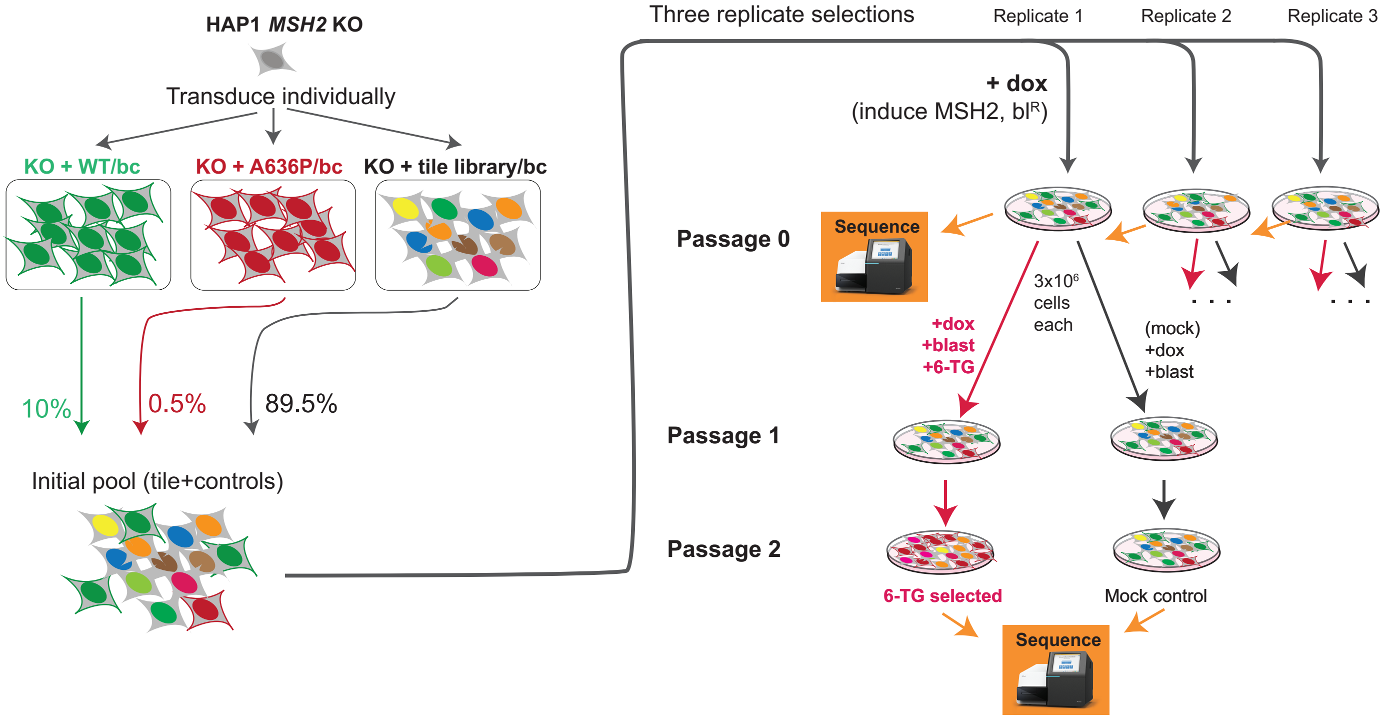


**Figure S6. Schematic of *MSH2* functional selection.**

Barcoded *MSH2* mutant cDNA libraries were stably integrated into HAP1 *MSH2* KO cells by lentiviral transduction. Cells bearing each *MSH2* mutant tile library (mixed colors) were combined with cells bearing WT (10%, green) or A636P (0.5%, red). Each initial pool was split into 3 replicates, expanded in the presence of doxycycline to induce *MSH2* expression (‘Passage 0’), then subjected to two rounds of mock or 6-TG selection, seeding $3x10^6$ cells per passage. Genomic DNA was harvested for *MSH2* tile-seq and barc-seq at passages 0 and 2. bc=barcode, $bl^R$=blasticidin resistance

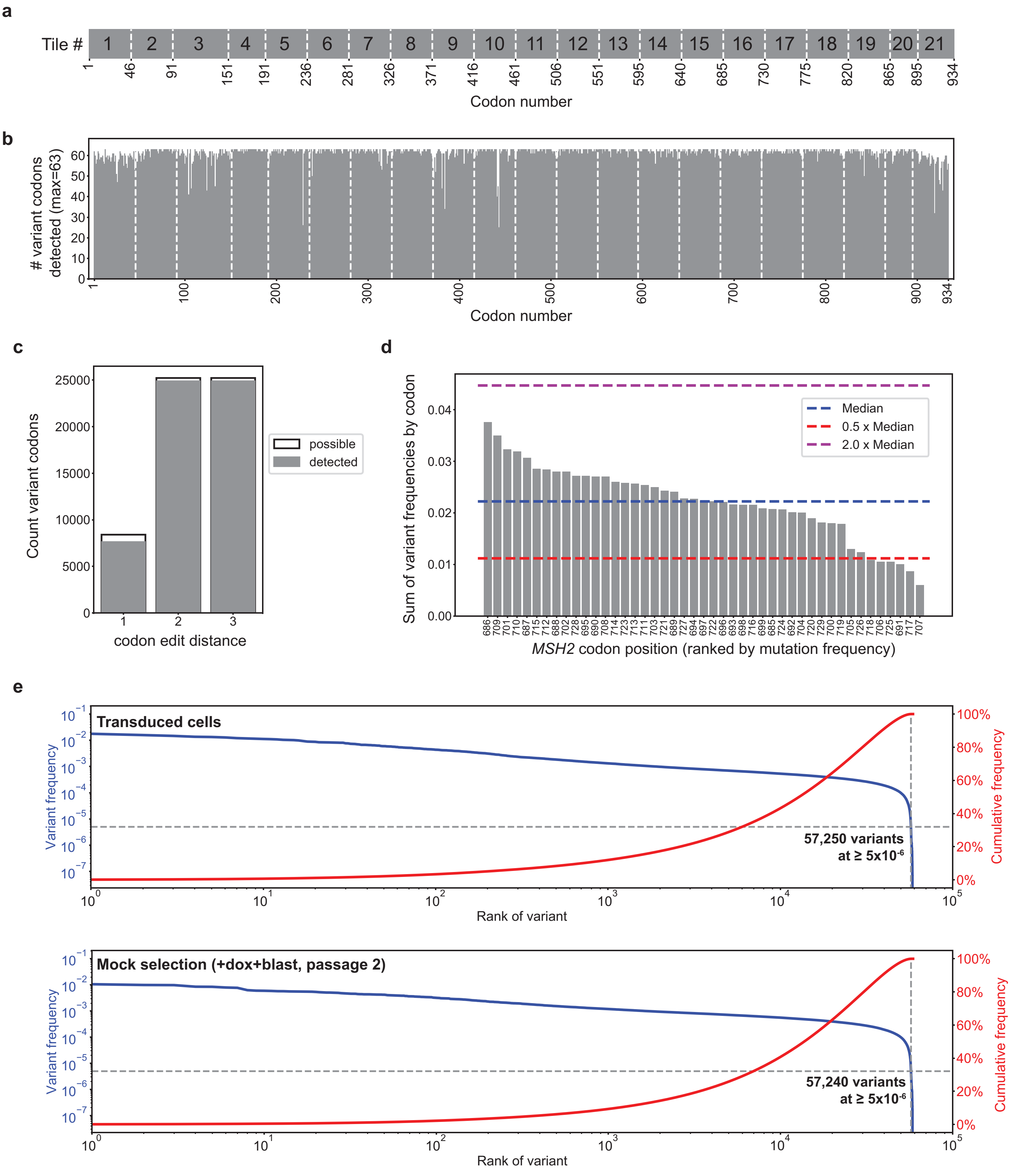


**Figure S7. Coverage and uniformity of integrated *MSH2* mutations.**

**a**, *MSH2* cDNA was divided into 21 tiles (dashed lines). Listed positions indicate the starting codon number of each tile. **b**, Coverage of possible mutations across *MSH2*. Plotted is the count of mutations observed (max.

possible=63), by codon position, following transduction (requiring per-mutation frequency $\geq 5 \times 10^{-6}$ after position-specific sequencing error correction to be considered present). Dashed white lines denote tile boundaries. **c**, Representation of *MSH2* mutations by edit distance. **d**, Uniformity of mutagenesis for a representative tile (tile 16), showing per-codon mutation frequency vs codon position (ranked by frequency). **e**, Waterfall plot of mutations' uniformity, with per-mutation frequency (blue; left y-axis) or cumulative frequency (red; right y-axis) plotted against mutation rank, for transduced cells (upper) and mock treatment (lower).

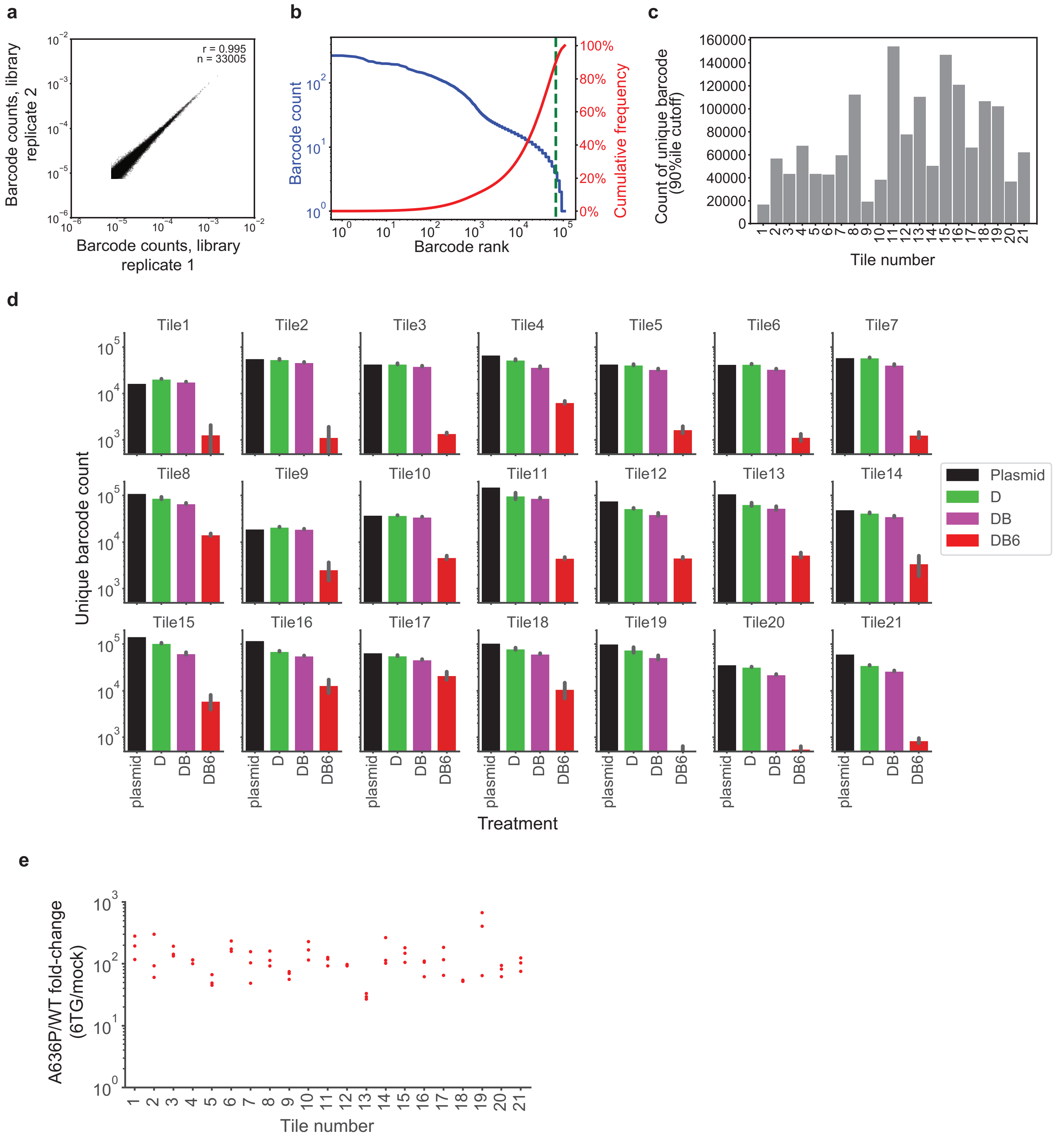


**Figure S8. Measuring internal control alleles throughout selection with barc-seq**

**a**, Technical reproducibility of barc-seq, comparing per-barcode normalized read counts of library replicates from two aliquots of the same gDNA (8 µg each). **b**, Quantifying the complexity of integrated *MSH2* library barcodes. Representative waterfall plot of integrated barcode abundances (tile 4), showing barcode frequency versus rank (blue curve; left y-axis) and cumulative frequency vs rank (red curve; right y-axis). The number of barcodes present was taken as the 90th percentile among read-ranked barcodes (dotted line), such that all barcodes with at least that many reads cumulatively account for 90% of read counts. **c**, Barcode complexity (at 90th percentile) of *MSH2* mutant plasmid libraries, by tile. **d**, Barcode complexity by stage of selection, per tile. Count of unique barcodes (90th percentile) is shown for each tile library, in the plasmid DNA (black), integrated in transduced cells (doxycycline-only; “D”, green), remaining after mock treatment (doxycycline+blasticidin only; “DB”, magenta) and post selection (doxycycline+blasticidin+6-TG; “DB6”, red). On average, each measured variant was associated by ≥8 barcodes. **e**, Enrichment of A636P control barcodes, relative to WT barcodes, in 6-TG versus mock selection.

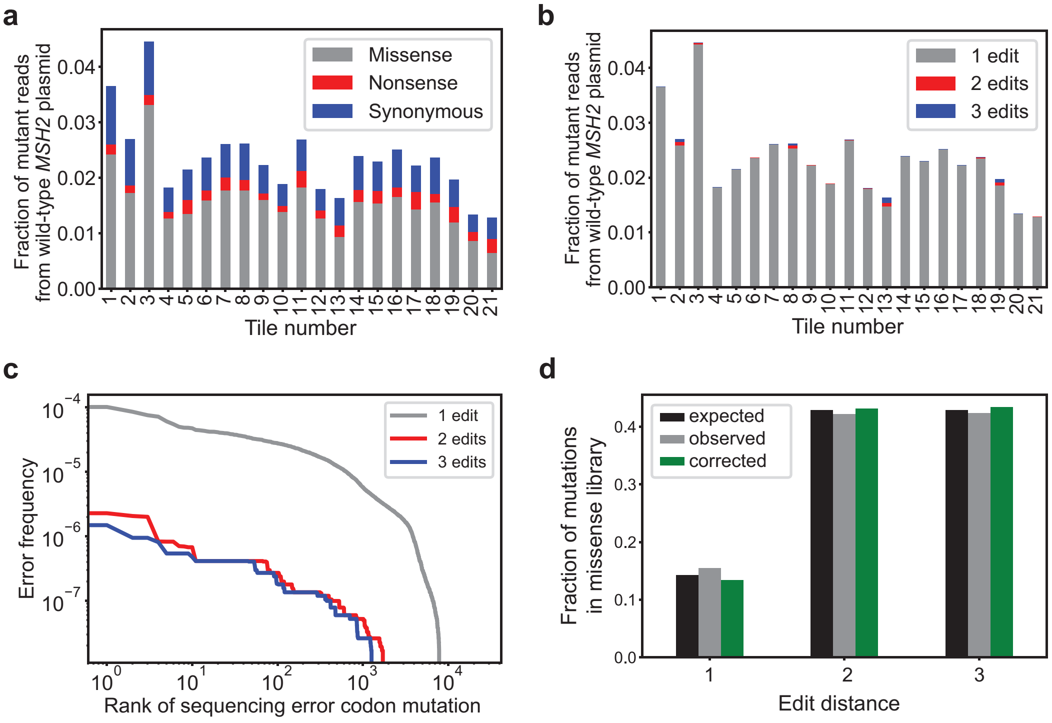


**Figure S9. Quantifying and compensating for sequencing errors.**

**a**, **b**, Per-codon sequencing error frequencies estimated by subjecting wild-type *MSH2* plasmid DNA to tile-seq analysis, summarized by tile number, and shaded by consequence **(a)** or number of base-pair edits **(b)**. Single-base pair, missense errors were the most common. **c**, Distribution of per-codon sequencing error frequencies versus rank, separated by the number of base pair edits (1 to 3), indicating a subset (~$10^3$) of single-base errors were relatively more frequent ($10^{-4}$ to $10^{-5}$). **d.** Within *MSH2* mutational libraries, the overall counts of codon mutations by edit distance (1 to 3) expected if perfectly random (black), those observed before correction (gray), and after subtraction of wild-type error frequencies (green), showing little change on aggregate from sequencing error correction.

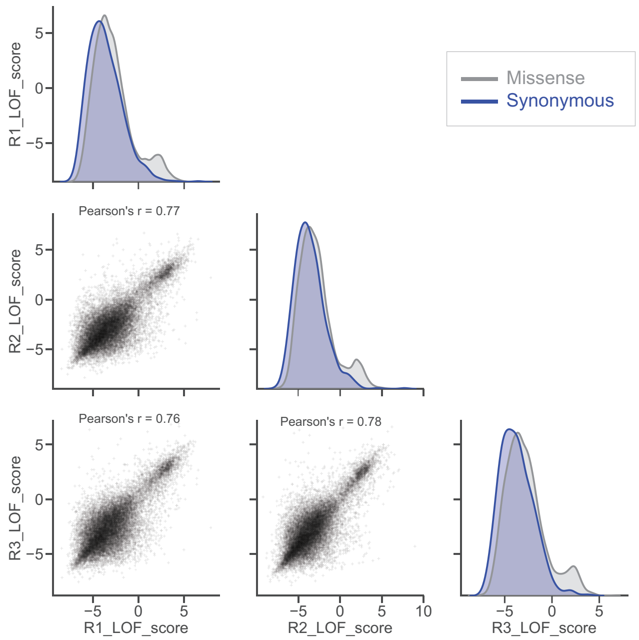


**Figure S10. Reproducibility across replicate selections.** Scatterplots comparing LOF scores for each pair of replicate selections; density plots show overall distribution of scores for missense (gray) and synonymous (blue) variants.

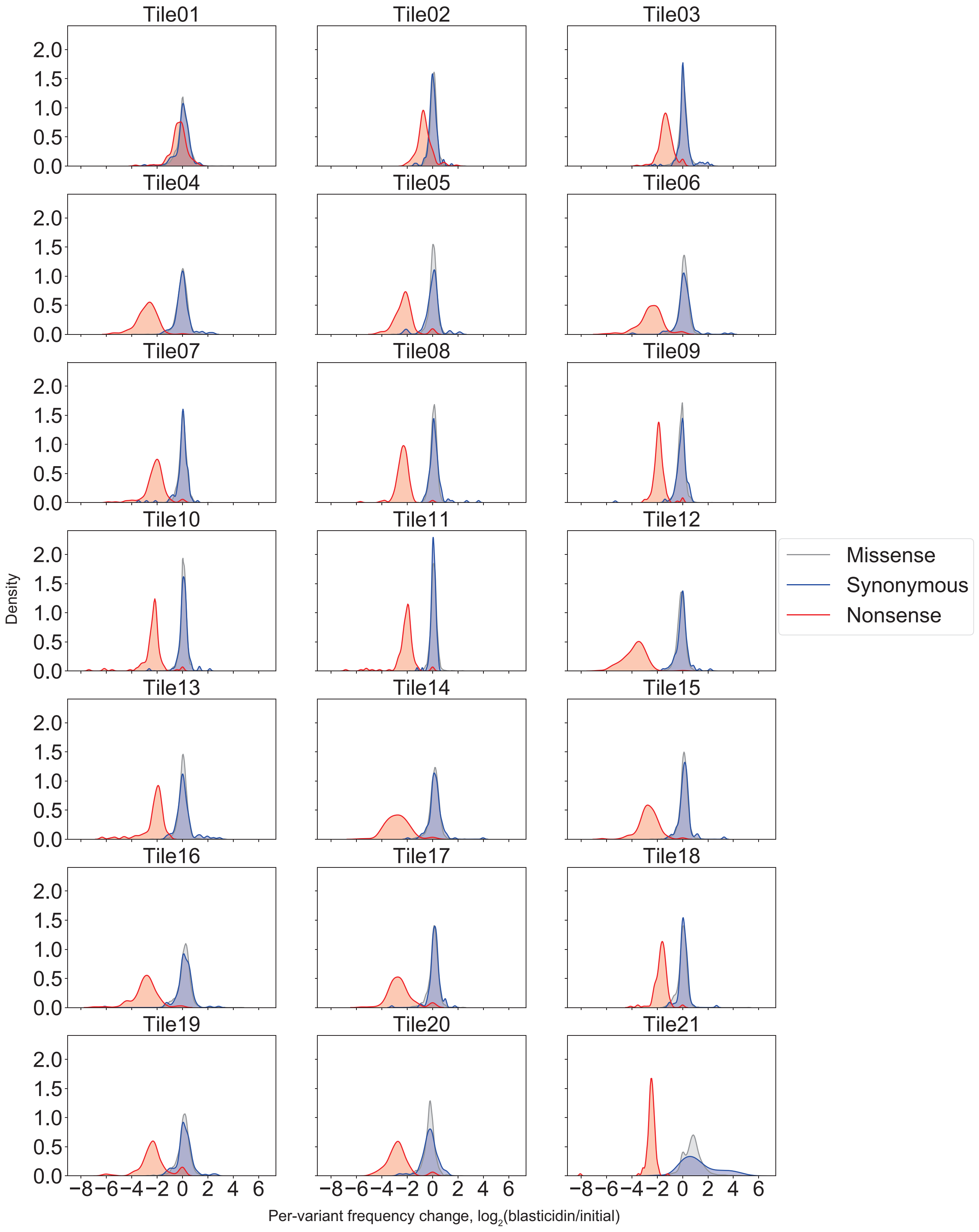


**Figure S11. Selection for full-length *MSH2* expression with in-frame blasticidin marker.**
Density plots showing change in variant frequency ($log_2$ ratio) after mock selection (blasticidin treatment) relative to the starting pool within each tile. Distributions are shaded by variant consequence: missense (gray), synonymous (blue), and nonsense (red). Blasticidin is selective against premature truncation in all tiles except tile 1, with comparatively little effect upon missense and synonymous variant clones; variants with log-ratio shifts outside of [-2,2] are shown here but were filtered out of the final score set (**Methods**).

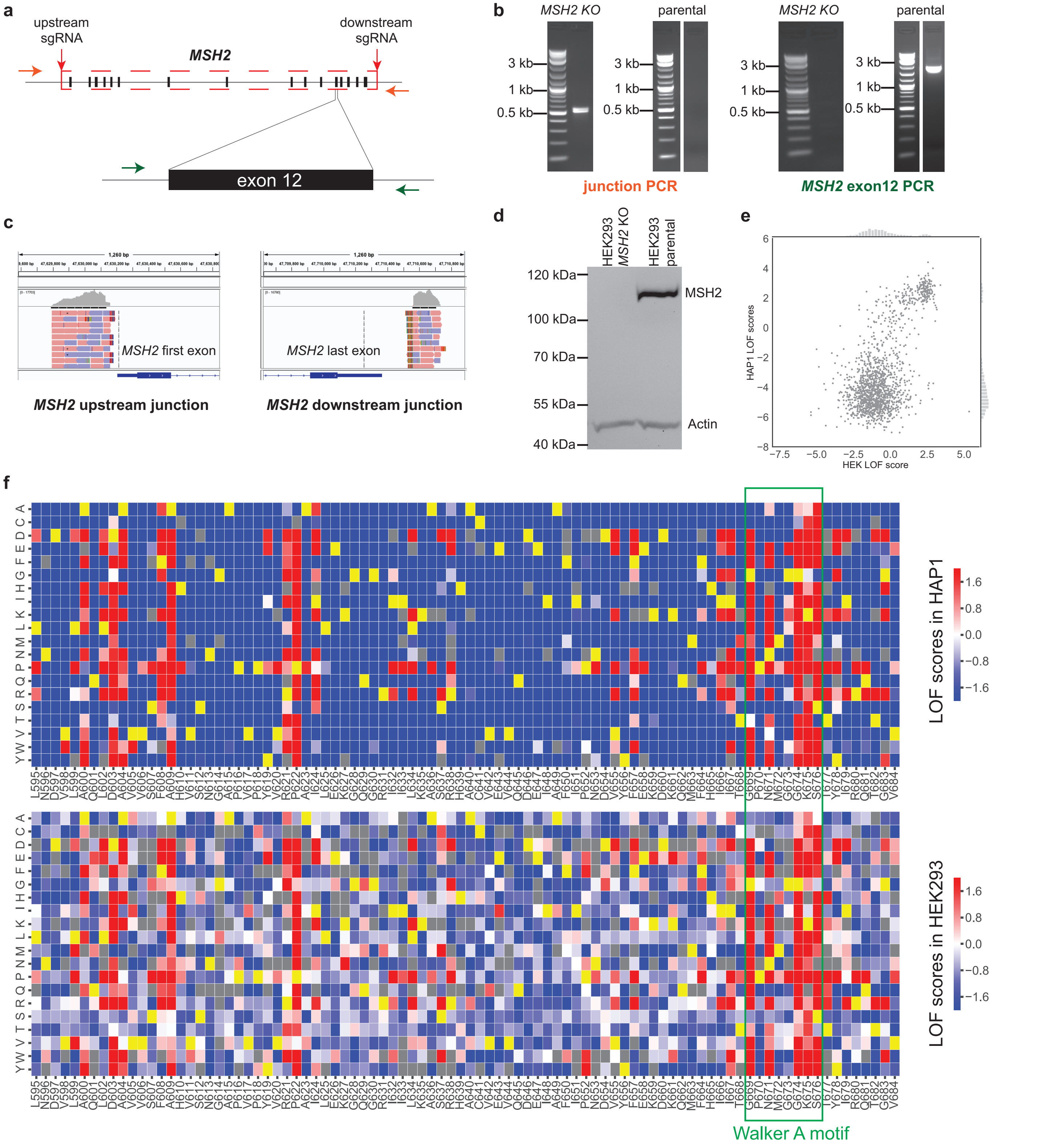


**Figure S12. Reproducibility in HEK293 cells**

**a**, *MSH2* whole-gene (80 kbp) deletion in HEK293 cells. **b**, Genotyping PCR detects gene deletion junctions only in knockout clone and internal product (exon 12) only in parental HEK293 cells. **c**, IGV screenshots showing upstream and downstream sides of deletion junctions. **d**, Western blot of MSH2 (and actin loading control) in HEK293 parental and HEK293 *MSH2* KO lines. **e**, Overall correlation between two cell lines' LOF scores for mutations in tiles 14 and 15. **f**, Sequence function heatmaps for LOF measured in HAP1 (upper) and HEK293 cell (lower), color scheme as in **Figure 2.** Green box denotes ATPase Walker A motif positions.

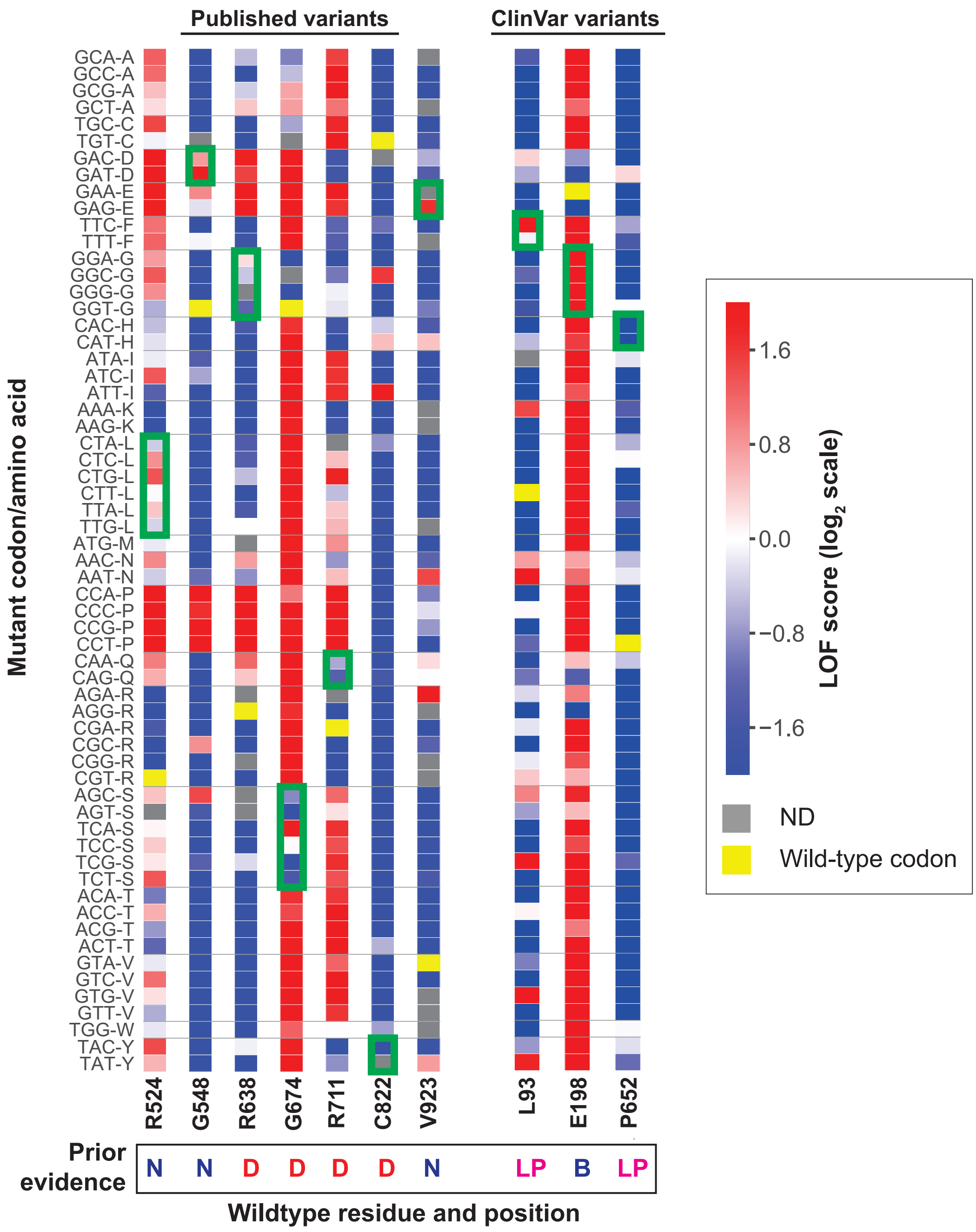


**Figure S13. Measurements discordant with prior evidence**

For missense variants with LOF scores discordant with existing functional data (left) or interpretations submitted to ClinVar (right), heatmaps showing LOF scores for each equivalent mutant codon contributing to the final, protein-level LOF score. Green boxes denote mutant codons which create the discordant missense mutation. D=deleterious, N=neutral; LP=likely pathogenic, B=benign, ND=no data.

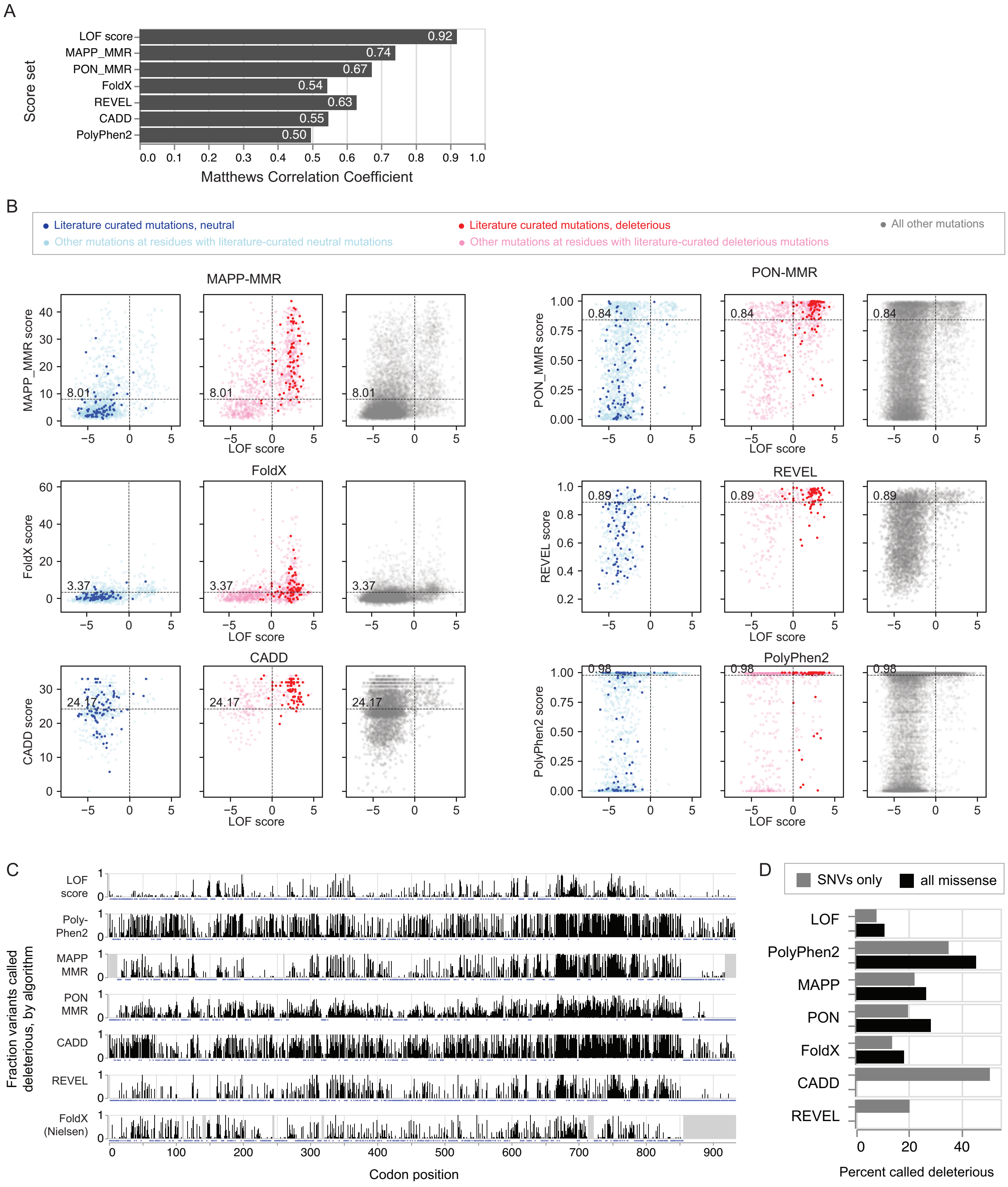


**Figure S14. Comparison with bioinformatic effect predictions.**

**a**, Classification performance (Matthews Correlation Coefficient, MCC) of experimentally measured LOF scores and bioinformatic predictions against published functional status of 126 *MSH2* missense mutations, at the optimal score threshold for each. **b**, Bioinformatic predictors' concordance (Jaccard Index) with experimental LOF scores is plotted versus algorithm-specific score threshold for deleteriousness. Algorithms were most concordant with LOF scores among mutations with published functional data (red), followed by other mutations at the same residues as those mutations (orange) and mutations all other residues (gray). **c**, Scatter plots of six algorithms' scores vs experimental LOF scores for literature curated mutations (neutral: blue, deleterious: red), other mutations at the same residues (light blue and pink, respectively), or all other mutations (gray). Label and horizontal lines denote score thresholds which maximized MCC on the training set. **d**, Distribution of loss of function fraction across *MSH2.* For experimental LOF scores and for each algorithm, the fraction of missense mutations at each residue called as deleterious is plotted by codon position. Gray box denotes no data; blue dots below baseline indicate positions where no variant was predicted to be deleterious. **e**, Overall percentage of missense mutations predicted to be deleterious by experiment (LOF) or algorithm, among all mutations (black; omitted for algorithms that score only single-nucleotide variants) or only those created by single-nucleotide variants (gray), at per-algorithm score cutoff used in **(a).**

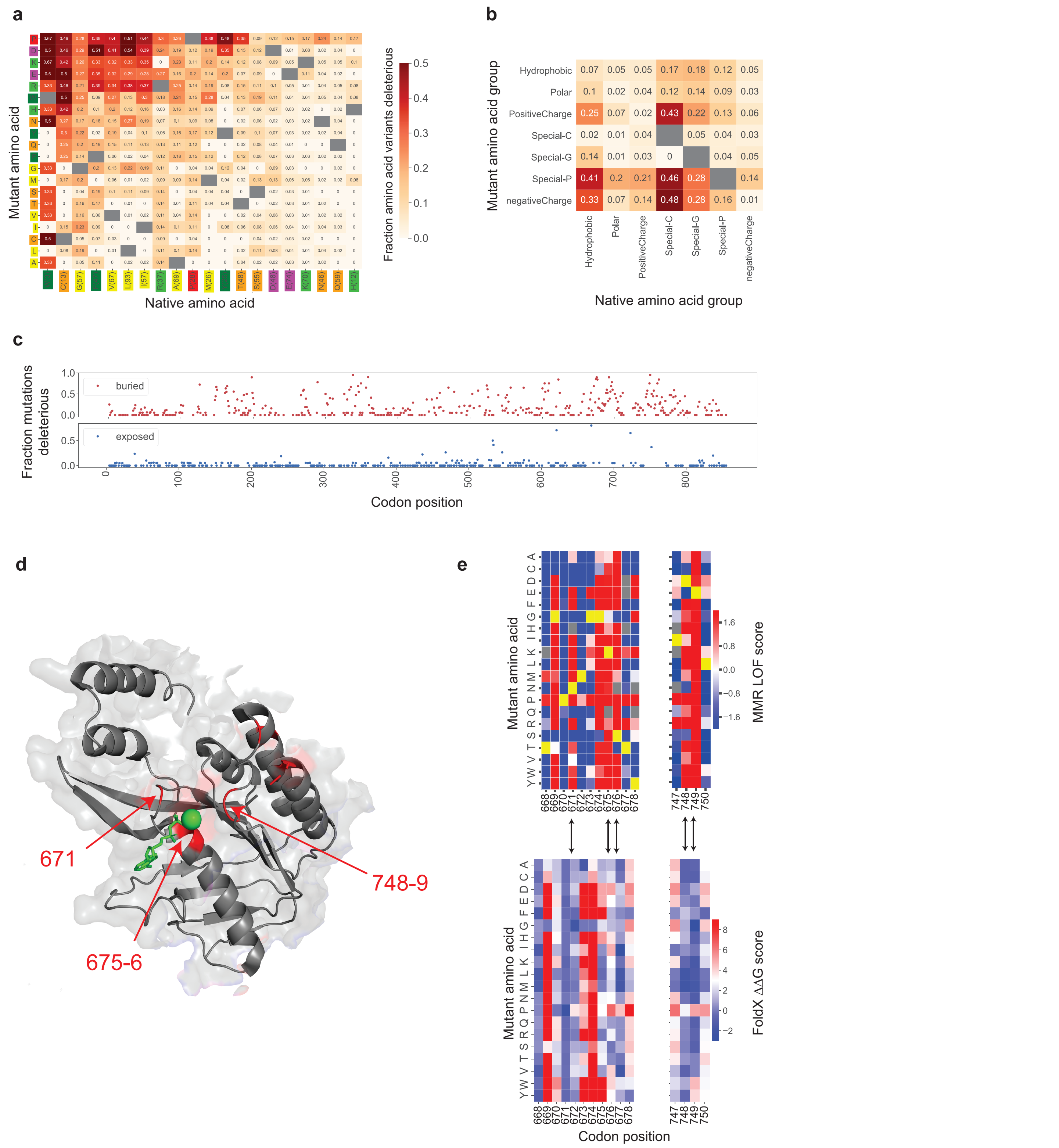


**Figure S15. Structural and chemical features of *MSH2* loss of function mutations.**
**a**, Deleterious fraction of mutations for each pair of amino acids. Amino acid symbols are shaded by chemical property (charge, size, etc); parentheses denote the number of each present in the native sequence. For instance, alanine occurs 69 times in MSH2; among those, replacement to proline was deleterious for 18/69 (0.26). Gray denotes no mutation (same amino acids). **b**, Same as **(a)**, but showing the deleterious fraction of mutations between chemically similar groups of amino acids (hydrophobic: V, Y, L, A, F, I, M, and W; polar: S, Q, T, and N; negatively charged: D and E; positively charged: R, K, and H; singletons: each of C, G, and P). **c**, Loss-of-function fraction plotted vs codon position for buried (red) and exposed (blue) residues. Buried residues

are less tolerant to substitutions (*P*<2.18e-42; two-sample KS test). **d.** ATPase domain from MSH2 crystal structure (PDB: 2O8E). Residues which are highly constrained (≥ 65% of mutations with LOF score > 0) but are not predicted to be destabilized (FoldX $\Delta\Delta G < 5$, (Nielsen et al. 2017)) are colored red, with residue numbers indicated. **e**, Heatmaps of LOF scores (top) and FoldX free folding energy $\Delta\Delta G$ (bottom) scores (Nielsen et al. 2017) at ATPase residues highlighted in (**d)**; arrows denote residues with strong constraint predicted by LOF score but not FoldX calculations.

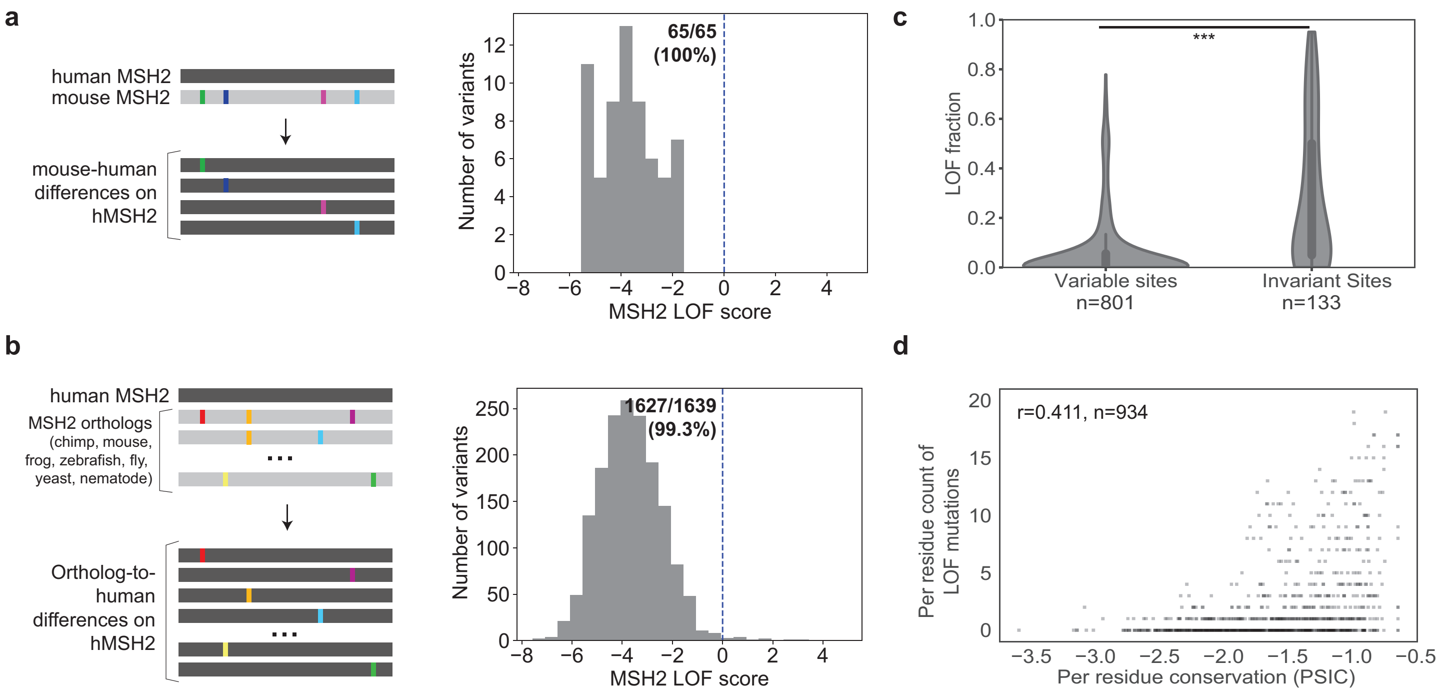


**Figure S16. Comparing LOF scores to divergent sites in *MSH2* orthologs**
**a**, Distribution of LOF scores for all point differences between the aligned human and mouse *MSH2* orthologs when individually present on the human sequence (inset: count and percent which are functionally neutral). Every mouse-human point difference was functionally neutral in the context of the human protein. **b**, Same as **(a)**, except using point differences to hMSH2 observed in a seven-way multiple alignment of eukaryotic MSH2 orthologs (*P. paniscus*, *M. musculus*, *X. tropicalis*, *D. rerio*, *S. cerevisiae*, *D. melanogaster*, *C. elegans*); nearly all (~99.3%) are neutral on hMSH2. **c**, Comparing fractions of possible mutations per residue which are loss-of-function, among residues which are variant among aligned eukaryotic MSH2 orthologs, or those which are invariant in those sequences (***, $P<1.7\times10^{-21}$, two-sample Kolmogorov-Smirnov test). **d.** Per-residue, count of possible mutations which are deleterious, versus conservation score (PSIC)
